# Multiresolution Clustering of Genomic Data

**DOI:** 10.1101/2025.06.13.659529

**Authors:** Ali Turfah, Xiaoquan Wen

## Abstract

Cluster analysis is a widely used unsupervised learning technique in genomic data analysis, with critical applications such as inferring genetic population structures and annotating cell types from single-cell RNA-seq data. However, most existing clustering methods focus on identifying a single optimal partition while overlooking intrinsic relationships among the inferred clusters. Moreover, clustering results produced by different algorithms often appear inconsistent, and there is a lack of principled approaches to extract shared, biologically meaningful patterns across diverse clustering outputs. In this work, we introduce a computational framework that enables systematic exploration of multi-resolution clustering structures in genomic data, starting from an initial configuration generated by *any* available clustering algorithm. The proposed algorithm provides a unified and principled approach for uncovering complex, nested biological relationships and reconciling discrepancies among clustering results. We demonstrate the utility of our framework through comprehensive simulations and applications to both genetic and single-cell transcriptomic datasets, highlighting its ability to recover interpretable and reproducible clustering structures. Furthermore, we show that our multi-resolution cluster analysis of complex genomic data yields valuable insights into patterns of human population migration and cell differentiation trajectories.

## 1 Introduction

Cluster analysis is a ubiquitous unsupervised machine learning approach to uncover latent structures and patterns in observed data. Clustering algorithms have been extensively applied to the analysis of genomic data, with important applications ranging from identifying population structures using densely genotyped genetic markers [1, 2, 3, 4, 5] to discovering novel cell types in single-cell RNA-seq (scRNA-seq) datasets [6, 7, 8, 9, 10]. These analyses play a crucial role in revealing biological insights from high-dimensional and complex genomic data.

Despite the availability of diverse clustering algorithms, interpreting clustering results remains a practical challenge in genomic data analysis [7]. Many existing methods and their benchmarking efforts overly emphasize identifying a flat partition and determining the “correct” number of clusters [10, 11, 12, 13, 14]. However, genomic data often exhibit nested or multi-scale structures that reflect underlying continuous biological differentiation processes. While a flat clustering may be deemed “optimal” under certain heuristic validation criteria or hyperparameter settings, it often overlooks the relative relationships among inferred clusters. For instance, in scRNA-seq analysis, focusing solely on a flat partition can obscure richer information about the relationships between cell types, failing to capture their relatedness arising from the cellular differentiation process [7, 15, 16]. We argue that a more appropriate approach is to represent the clustering of biological data hierarchically, allowing for the characterization and quantification of relationships between clusters on a universally interpretable scale. These relationships can be conceptualized as a continuum of resolutions or “zoom levels,” where lower resolutions (zoomed out) highlight broad distinctions and higher resolutions (zoomed in) reveal finer-grained differences.

Another challenge lies in reconciling the differences in results produced by diverse clustering algorithms. These differences often stem from the varying operational definitions of “clusters” employed by different methods [17, 18]. While all methods generally agree that clusters should exhibit internal cohesion (i.e., observations within a cluster are similar to each other) and external isolation (i.e., observations in different clusters are disssimilar to each other) [19, 20, 21], there is no universally accepted metric for quantifying these properties, and different clustering methods formalize and implement these criteria in distinct ways. For example, community detection methods (e.g., Louvain and Leiden algorithms) rely on measures of graph connectivity [22, 23], density-based methods (e.g., DBSCAN and DensityCut) identify clusters based on peaks and valleys in estimated data density [24, 25, 26], and centroid-based methods (e.g., *k*-means) aim to minimize within-cluster pairwise distances. Because these operational definitions are not fully compatible, it is unsurprising that in practice they can yield markedly different clustering results [27]. Nevertheless, identifying common patterns across diverse clustering results can be highly beneficial, as consensus patterns emerging from distinct procedures are more likely to reflect genuine biological signals rather than artifacts of any specific algorithm. This aligns with the principle of inferential reproducibility [28], which emphasizes the ability to draw consistent conclusions from different analytic choices applied to the same dataset. In the context of cluster analysis, leveraging agreement among outcomes from diverse algorithms offers a principled way to enhance the reliability and interpretability of the inferred structures. However, despite its conceptual appeal, analytical implementations of this principle are still lacking in current clustering practice.

In this paper, we address the above challenges by proposing a computational framework for multi-resolution cluster analysis of genomic data. Specifically, we introduce two key technical innovations. First, we define a probabilistic measure, *P*_mc_, grounded in statistical decision theory to quantify the overall level of external isolation relative to internal cohesion within a given clustering configuration. This metric is motivated by a natural intuition: if clusters are well separated, the original cluster membership of any data point should be easily recoverable with high confidence. Second, building on the appealing mathematical properties of this measure, we develop the *P*_mc_ hierarchical merging (PHM) algorithm and accompanying visualization tools, which enable systematic exploration of clustering structures across multiple resolutions from any initial clustering configuration. Importantly, both *P*_mc_ and the PHM algorithm are broadly applicable to any existing clustering method. Consequently, our framework offers a unique advantage in identifying consensus cluster patterns across different resolutions and outcomes generated by different algorithms.

In the remainder of the paper, we describe this multi-resolution clustering framework in detail and demonstrate its utility through applications to both synthetic datasets and real genomic data, including examples from human population genetics and single-cell RNA-seq analysis. The software package implementing the proposed PHM algorithm is freely available at https://github.com/aturfah/phm.

## 2 Results

### 2.1 Method Overview

#### 2.1.1 Probabilistic Quantification of Internal Cohesion and External Separation

The foundation of our proposed computational method is the probabilistic quantification of external separation with relative to internal cohesion for a given clustering configuration. We first interpret each inferred cluster as representing an underlying population, with its internal cohesion fully characterized by the corresponding population distribution, *P* (***x*** | *θ*), where ***x*** denotes the data measurement and *θ* denotes the population or cluster label. For clustering methods that do not directly estimate *P* (***x*** | *θ*) but instead partition the observed samples, we estimate the underlying population distributions corresponding to these partitions using Gaussian mixture models – a well-known universal approximator of arbitrary densities [29]. Next, we introduce a novel probability-based metric, *P*_mc_, to quantify the external separation of all inferred clusters, which we refer to as the distinguishability criterion. *P*_mc_ is defined as the probability of failing to assign a data point to its generating cluster, averaged over all possible data points. Intuitively, *P*_mc_ *→* 0 indicates a high degree of separation among inferred clusters, as the probability of erroneously assigning any data point to an incorrect cluster is minimal. Conversely, substantial overlap between any clusters leads to a significant increase in *P*_mc_.

*P*_mc_ is derived within a rigorous decision-theoretic framework and has several useful properties that can be leveraged to assess and validate inferred clustering structures in traditional cluster analysis (Methods). Here, we focus on its most relevant property for the proposed multi-resolution clustering algorithm – the merging property. Notably, merging any two existing clusters into a single one always decreases *P*_mc_ (Proposition 1 in Methods). We define 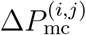 as the reduction in *P*_mc_ when a pair of clusters, *i* and *j*, are merged. The value of 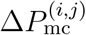 captures the relative distance between clusters *i* and *j* on a probability scale: if they are well separated, merging leads to a minimal decrease in the overall misclassification probability; conversely, if they significantly overlap, the reduction in *P*_mc_ is substantial. Moreover, *P*_mc_ can be expressed as the sum of all pairwise Δ*P*_mc_ values of a given clustering configuration, i.e.,

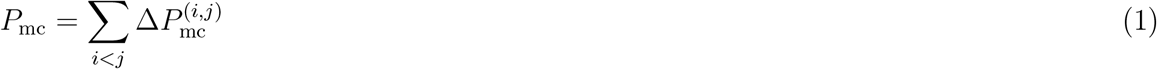

#### 2.1.2 PHM Algorithm

The properties of Δ*P*_mc_ and Equation (1) motivate an intuitive and straightforward algorithm for exploring clustering structures at different resolutions. We refer to this algorithm as the *P*_mc_ Hierarchical Merging (PHM) algorithm.

Starting from the finest resolution – the optimal clustering configuration determined by a given clustering algorithm – the PHM algorithm iteratively merges the two closest clusters in the current configuration based on their Δ*P*_mc_ values. This process continues until only a single cluster remains, at which point *P*_mc_ = 0. The detailed procedure of the PHM algorithm, including the dynamic updating of 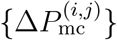 and *P*_mc_ values, is provided in Algorithm 1 in Methods. By iteratively merging clusters, the PHM algorithm systematically transitions from a finer resolution clustering to increasingly coarser resolutions, forming progressively agglomerated clusters at each step.

Figure 1 illustrates the PHM algorithm applied to a simulated dataset. The initial configuration consists of five clusters inferred using a Gaussian mixture model (GMM). The merging process in the PHM algorithm is visualized through a dendrogram, which highlights the order in which cluster pairs are merged. For a subtree 𝒮, consisting of *m* leaf nodes (i.e., original clusters) and formed by directly merging two smaller subtrees, 𝒮_*l*_ and 𝒮_*r*_, we define its depth as a decreasing function of 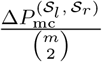, which intuitively represents the average pairwise similarity among the original clusters within 𝒮. Thus, a greater subtree depth indicates the inclusion of more distantly separated (i.e. more diverse) original clusters.

**Figure 1:**
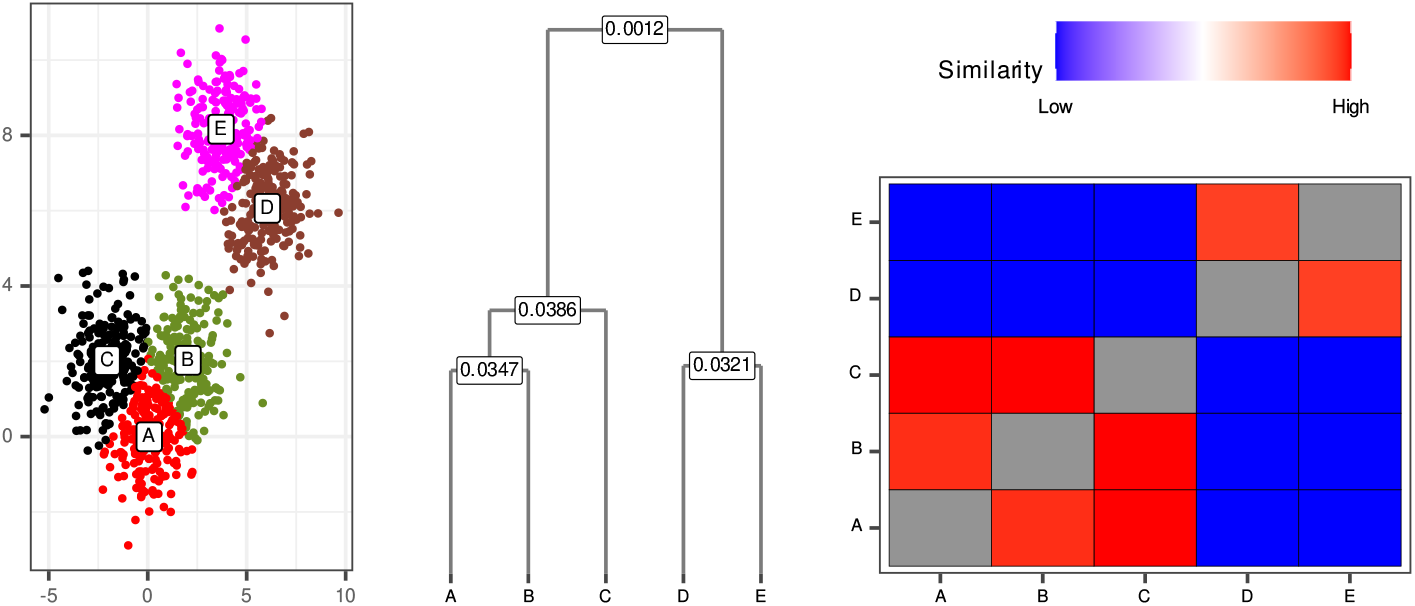
Visualization of the PHM procedure. *(Left)* Scatterplot of the simulated data colored by the true cluster labels (A - E). *(Center)* Dendrogram illustrating the PHM merging procedure, where the value at each internal node represents the corresponding Δ*P*_mc_ of the merge. *(Right)* Heatmap visualizing the pairwise similarity between the 5 initial clusters, with color intensity reflecting the *ρ* value between a pair of initial clusters. The pattern of the overall structure of the clusters can be read from the the dendrogram as well as the heatmap.

The varying cluster structures at different resolutions can be interpreted from the dendrogram by observing changes in the scale of subtree depths. In the illustrative example, the dendrogram shows that the original five clusters form two distinct agglomerated clusters at a coarser resolution, indicating that original clusters (*A, B, C*) are more closely related to each other, while clusters (*D, E*) are also more closely grouped together.

Additionally, we provide a heatmap representing the pairwise relationships among all original clusters. The pairwise measure between clusters *i* and *j*, denoted as *ρ*(*i, j*), corresponds to the Δ*P*_mc_ value at which clusters *i* and *j* are first merged into the same agglomerated cluster during the merging process. Specifically,

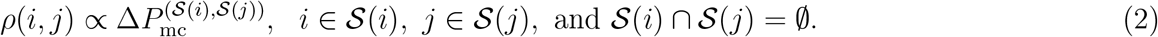

Closely related clusters tend to have higher *ρ* values, while those that are more distant have *ρ* values approaching zero. The block structure revealed in the heatmap (Figure 1) effectively captures the complex clustering structures at different resolutions.

### 2.2 Synthetic Data Examples

Additional details on the simulation studies below can be found in Supplementary Methods 1.

#### 2.2.1 Nested Diamonds

Our first numerical experiment highlights the importance of recognizing varying cluster structures across different resolutions. At the finest resolution, we simulate 100 observations from each of 64 distinct Gaussian distributions in a 2D plane (Figure 2a). The simulated data are further arranged as follows to form a complex pattern: (i) each set of four Gaussian clusters forms a small diamond (inner diamond), resulting in 16 inner diamonds; (ii) each set of four inner diamonds forms a larger diamond (outer diamond), yielding four outer diamonds; (iii) the four outer diamonds are positioned along the edges of a square. As a result, no single clustering structure at any fixed resolution can fully capture the complex, multi-scale relationships among the subgroups.

**Figure 2:**
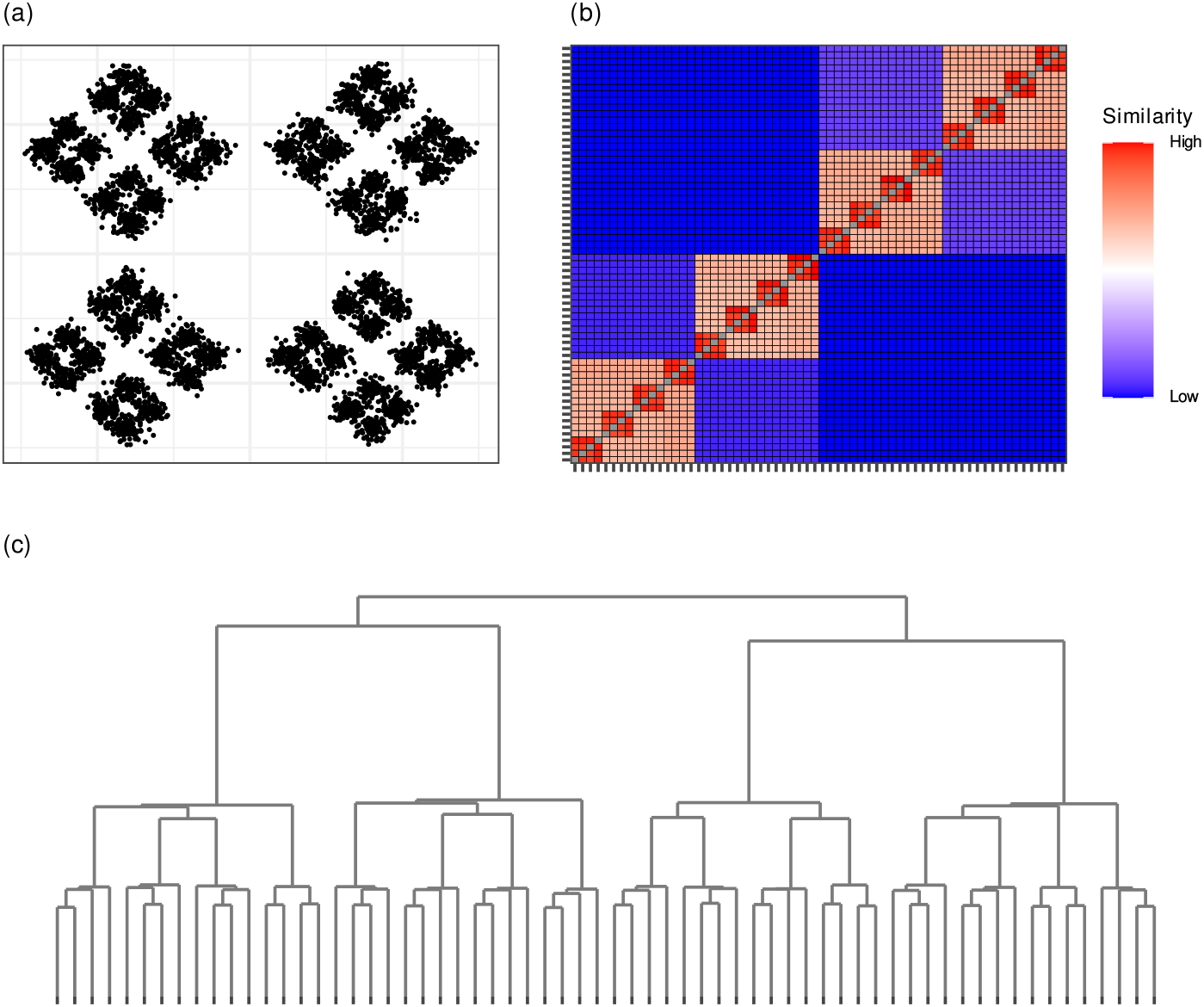
PHM detects multi-level clustering structures in the nested diamonds. (a) Simulated data arranged in a nested diamond pattern. Clusters are organized into three hierarchical layers: 16 inner diamonds, 4 outer diamonds (each formed by grouping four inner diamonds), and a single outer square composed of the four outer diamonds. (b) Heatmap of Δ*P*_mc_ values along the branches of the PHM dendrogram initialized from the GMM solution, revealing three distinct levels of cluster structure. Dark red regions capture strong similarity within inner diamonds, orange regions correspond to outer diamonds, and dark blue regions represent the grouping of outer diamonds along the edges of the square. Groupings of similar color indicate structures occurring at the same resolution. (c) PHM dendrogram derived from the GMM clustering result. Subtree heights cluster at three levels, corresponding to the nested hierarchy in the data: bottom-level merges reflect the formation of inner diamonds, mid-level merges form outer diamonds, and top-level merges correspond to the final aggregation into the square.

With appropriate hyperparameter values and initialization, commonly used clustering algorithms – including *k*-means, GMM, hierarchical clustering, and DensityCut – correctly identify the 64 clusters at the finest resolution. We then apply the PHM algorithm using these initial clustering results as input. Figures 2b and 2c illustrate the PHM output based on the GMM results. The resulting dendrogram and heatmap indicate that the PHM algorithm accurately captures the three levels of nested structure in the data. The PHM results for the other clustering methods are consistent with the GMM results (Supplementary Figures 1-3).

We note that HDBSCAN [30, 31], a clustering method that extends the popular density-based algorithm DBSCAN, and hierarchical clustering produce a dendrogram similar to Figure 2b when applied to the simulated dataset (Supplementary Figures 4-5). Nevertheless, compared to HDBSCAN, the PHM algorithm does not restrict the choice of clustering method and can be applied to the outputs of any clustering algorithm. The observation that PHM yields consistent results across initial configurations generated by a variety of clustering algorithms highlights its general applicability and robustness.

#### 2.2.2 Double Ring

In this simulation study, we examine PHM’s ability to reconcile clustering results from different algorithms and to identify common underlying structures. The simulated dataset comprises 5,000 two-dimensional points randomly drawn from two nested rings (Figure 3), with points uniformly distributed within radii 2–3 for the inner ring and 6–7 for the outer ring. We perform cluster analysis using DensityCut (density-based), GMM (model-based), *k*-means (centroid-based), hierarchical clustering (with Ward linkage, connectivity-based), and the Louvain algorithm (community detection-based).

**Figure 3:**
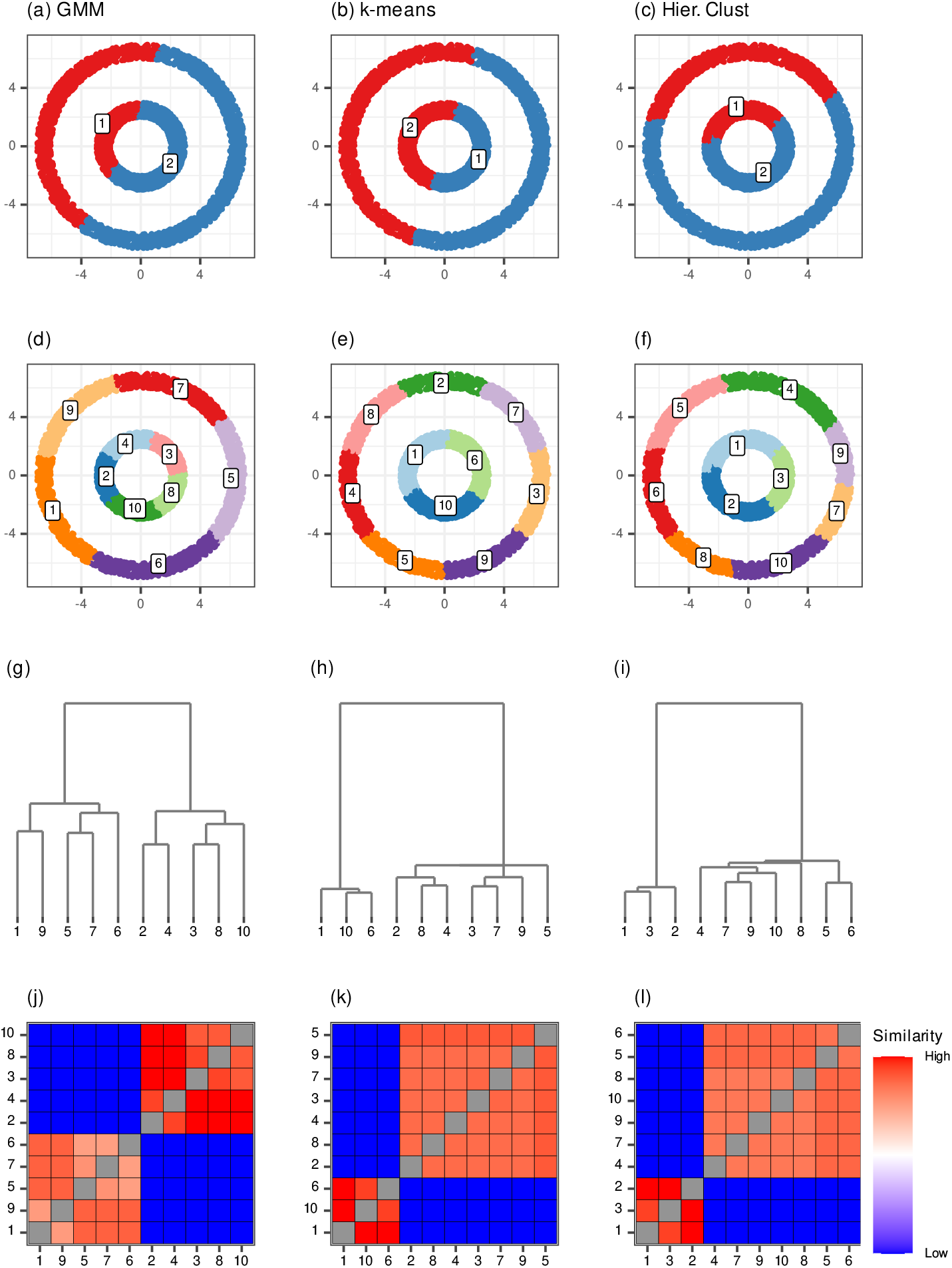
PHM recovers the double-ring structure from the outputs of diverse clustering algorithms. Results for the double-ring dataset using GMM, *k*-means, and hierarchical clustering (Hier. Clust) are shown in columns from left to right. *(a–c)*: Partitions obtained by explicitly setting *K* = 2. *(d–f)*: Optimal partitions selected by each algorithm based on its respective model selection criterion. *(g–i)*: PHM dendrograms constructed from the corresponding optimal solutions of GMM, *k*-means, and hierarchical clustering. *(j–l)*: Heatmaps visualizing PHM results for the same initial configurations. Together, the dendrograms and heatmaps demonstrate that PHM consistently recovers the underlying double-ring structure across all clustering inputs.

Because the two rings are not linearly separable, the dataset presents a significant challenge for many popular clustering algorithms attempting to recover the inner and outer rings as distinct clusters. Under their respective default settings, GMM, *k*-means, hierarchical clustering, and the Louvain algorithm all over-segment the data, dividing each ring into multiple subgroups (Figure 3 e-f, Supplementary Figure 6). Moreover, even when explicitly instructed to identify *two* clusters, GMM, *k*-means, and hierarchical clustering fail to recognize the true double ring structure (Figure 3 a-c). In contrast, only DensityCut and the Louvain algorithm (with carefully tuned hyperparameters) can correctly identify the nested rings as two clusters (Supplementary Figures 6-7). This example highlights the lack of inferential reproducibility among popular clustering methods and the challenges of reconciling their results.

We find that the PHM algorithm is particularly effective at revealing the shared structure across seemingly inconsistent clustering results: starting from each algorithm’s optimal clustering configuration under their respective default settings, PHM consistently recovers the true double-ring structure, as illustrated by both the dendrograms and the heatmaps (Figure 3 g-l, Supplementary Figure 6). More broadly, this simple experiment illustrates that higher resolution structures uniquely identified by a single algorithm may be driven by unverified, algorithm-specific modeling assumptions, whereas structures consistently recovered across multiple algorithms are more likely to reflect real patterns.

### 2.3 Population Structure Inference from Human Genome Diversity Project Data

In this illustration, we apply the PHM algorithm to cluster genetic data from the Human Genome Diversity Project (HGDP) [32, 33], with the goal of identifying population structure. The dataset includes 927 unrelated individuals sampled globally and genotyped at 2,543 autosomal SNPs. Geographic sampling locations are broadly categorized into 7 continental groups: Europe, Central/South Asia (C/S Asia), Africa, the Middle East, the Americas, East Asia, and Oceania. Following standard procedures, we preprocess the genotype matrix using principal component analysis (PCA) and select the first 9 principal components (PCs) based on the elbow plot.

Figure 4 presents the PHM output based on the initial clustering from the GMM algorithm. The optimal GMM solution, selected by BIC, identifies six mixture components/clusters. Individuals from Africa, the Americas, East Asia, and Oceania each correspond largely to distinct clusters. The remaining two clusters represent individuals from Central/South Asia, Europe, and the Middle East, reflecting shared genetic ancestry and admixture among these groups. The relative distances among clusters can be inferred from the PHM merging order and its visualization. For instance, the first merge joins clusters primarily comprising samples from Europe, the Middle East, and Central/South Asia, suggesting that these three populations share the closest genetic distances. Notably, the relative genetic relationships among GMM clusters, as inferred by PHM (shown in the dendrogram and heatmap), roughly align with – and can be interpreted by – the likely path of historical human migrations: from Africa into the Middle East; from the Middle East to Europe and Central/South Asia; from Central/South Asia to East Asia; and from East Asia to Oceania and the Americas. Analyses using haplotype information [33] and high-coverage genome sequencing data [34] also support this finding.

**Figure 4:**
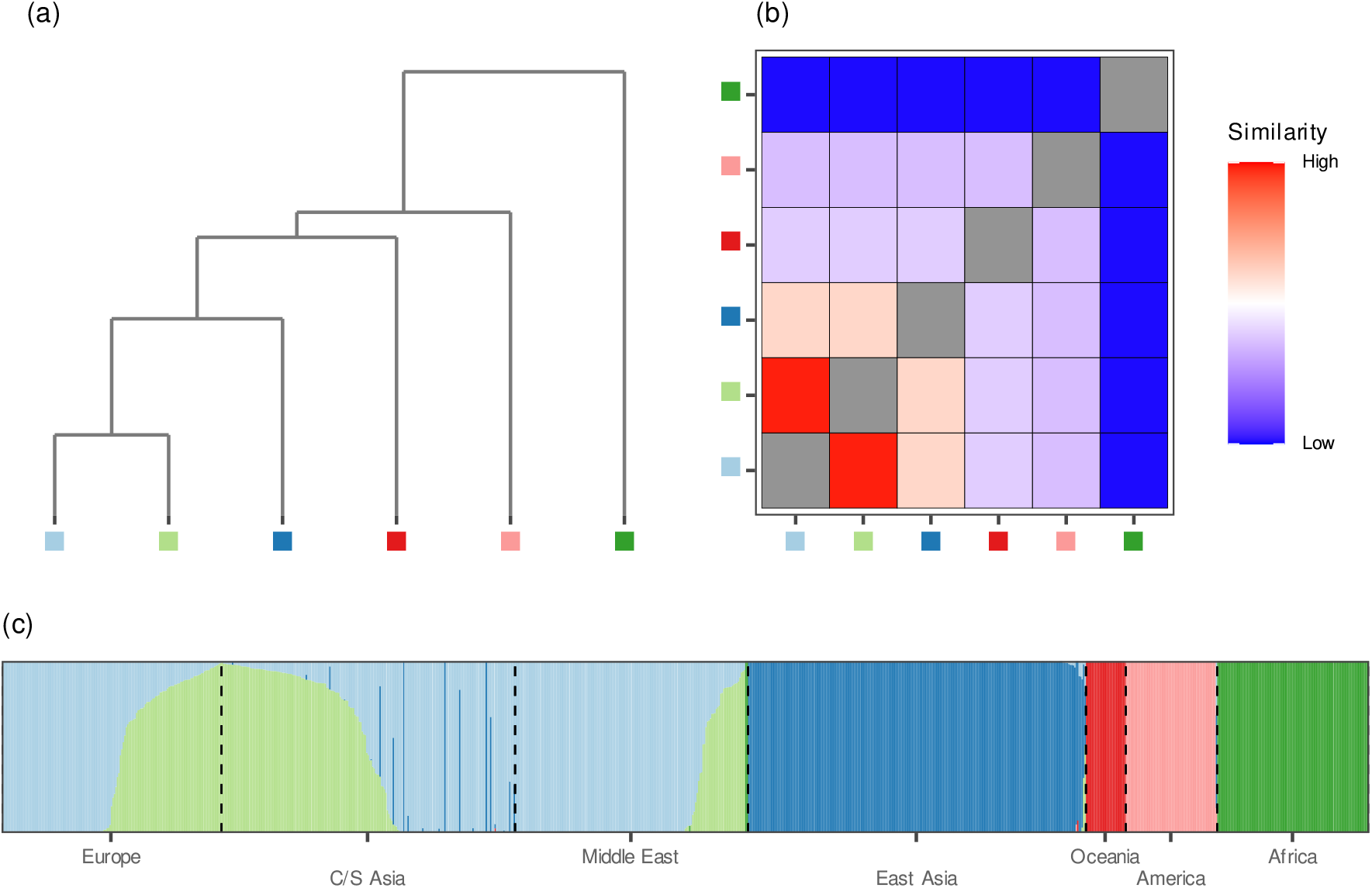
PHM result from HGDP data analysis. The PHM results are based on the output from the Gaussian mixture model (GMM). (a) PHM dendrogram illustrating the hierarchical merging of 6 estimated mixture components. (b) PHM heatmap showing the pairwise similarity between the initial clusters. (c) Distruct plot [35] of posterior component assignment probabilities for the 927 HGDP samples, with individuals grouped by geographic sampling location and colors indicating mixture components. The relationships among clusters revealed by the PHM dendrogram and heatmap reflect the genetic distances between major population groups.

As alternatives to the GMM, we also evaluate DensityCut, *k*-means, hierarchical clustering (using Ward linkage), and the Leiden algorithm to generate initial clustering results. The optimal clustering configurations, selected based on each method’s respective optimality criterion, vary in resolution (Supplementary Figures 8-11). All methods except the Leiden algorithm yield coarser partitions with 4 to 5 clusters, often grouping individuals from Central/South Asia, Europe, and the Middle East into a single cluster. The Leiden algorithm produces a clustering configuration broadly similar to the GMM, but further divides the mixed samples from Europe, Central/South Asia (C/S Asia), and Africa into two distinct partitions. The PHM outputs based on these initial clustering configurations are largely consistent, suggesting that different algorithms characterize the relative genetic distances between major population groups similarly.

Finally, we repeat the analysis using HDBSCAN on the same PC-processed input data. The results appear to be strongly influenced by a small group of Oceania samples and are considerably less interpretable than the rest of the PHM results (Supplementary Figure 12, Supplementary Table 1).

### 2.4 Cluster Analysis of Human Peripheral Blood Mononuclear Cell Data

In this analysis, we apply the PHM algorithm to single-cell RNA sequencing (scRNA-seq) data from 2,638 human peripheral blood mononuclear cells (PBMCs) [36] to compare consensus structures obtained from different clustering approaches. Expression data for 13,714 genes, sequenced on the Illumina NextSeq 500, are normalized and scaled using the preprocessing procedures implemented in the Seurat package following the standard quality control. We perform principal component analysis (PCA) and select the first 9 components based on the elbow plot. Each cell is annotated as one of the following types using known biomarkers: naïve CD4+ T cells, memory CD4+ T cells, CD8+ T cells, natural killer (NK) T cells, B cells, CD14+ monocytes, FCGR3A+ monocytes, dendritic cells (DCs), or platelets. These cell type annotations are used for external validation but are not incorporated into the cluster analysis.

We perform cluster analysis using Gaussian mixture models (GMM), *k*-means, hierarchical clustering, DensityCut, and the Leiden algorithm. The optimal clustering configurations, selected according to each method’s respective criterion, exhibit substantial variability (Figure 5). The coarsest solution, produced by hierarchical clustering using the Silhouette score, identifies only two clusters: one comprising B cells and various T cell subtypes, and the other consisting of monocytes and dendritic cells. These two clusters broadly align with lymphoid and myeloid progenitor lineages, respectively. The solutions from the other clustering algorithms divide the cells into finer subgroups. The partition generated by the Leiden algorithm identifies four distinct groups that roughly correspond to T cells, B cells, monocytes, and dendritic cells (with platelets). The optimal *k*-means solution yields five clusters, further distinguishing NK T cells from other T cell populations. The GMM solution identifies seven clusters, separating CD4+ T cells from CD8+ T cells and distinguishing CD14+ monocytes from FCGR3A+ monocytes. Finally, DensityCut identifies ten groups, in which naïve and memory CD4+ T cells are split, and B cells and CD14+ monocytes are each divided into two distinct clusters.

**Figure 5:**
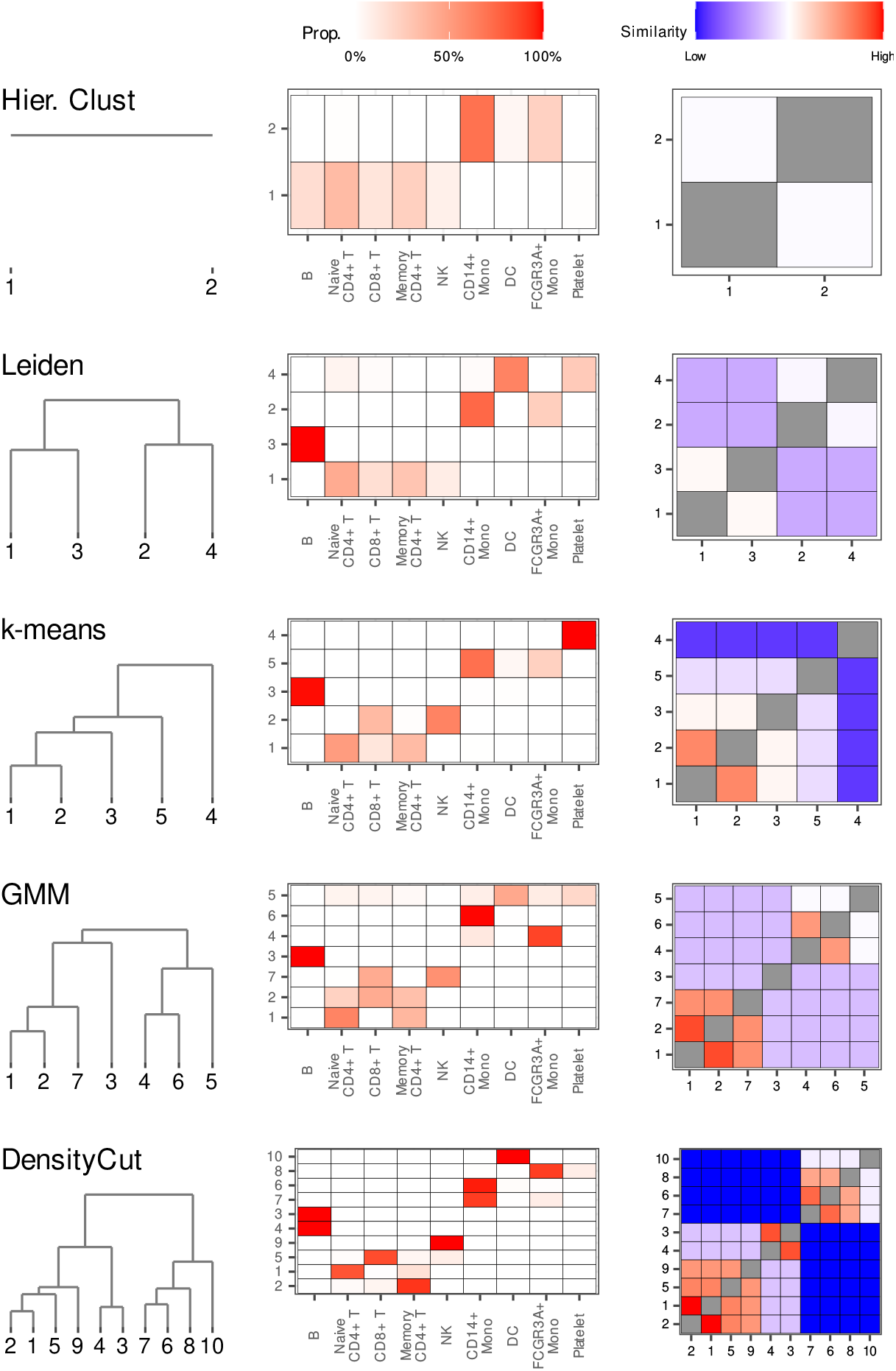
Cluster analysis of the PBMC data. Clustering results, along with the corresponding PHM dendrograms and heatmaps, are shown for the hierarchical clustering, Leiden, *k*-means, GMM, and DensityCut. Results for each clustering method are presented in rows. Columns are organized as follows: *(Left)* PHM dendrogram initialized from the clustering output. *(Center)* Cell-type composition of each inferred initial cluster, with color indicating the proportion of cells from each annotated cell type. *(Right)* PHM heatmap visualizing the similarity between pairs of initial clusters.

Running the PHM algorithm on these markedly different configurations (Figure 5) confirms that the clustering configurations reported by different algorithms correspond broadly to cell type classifications at varying resolutions, with finer-grained distinctions emerging at higher resolutions. Importantly, the PHM algorithm consistently recovers lower-resolution structures within the higher-resolution solutions across different initial configurations. The PHM results also reveal that T cell and B cell clusters are more closely related to each other, as are the monocyte and dendritic cell clusters. Overall, the dissimilarity patterns among clusters revealed by PHM align well with established immune cell differentiation pathways: T and B cells both derive from lymphoid progenitor cells, while monocytes and dendritic cells originate from myeloid progenitor cells. These findings indicate that the PHM algorithm effectively captures key features of the underlying cell differentiation trajectories from cluster analyses.

### 2.5 Cluster Analysis of Larval *C. elegans* Data

In our final illustration, we apply the PHM algorithm to a sample of scRNA-seq data from larval *C. elegans* cells [37]. The dataset contains expression counts for 20,271 genes across 32,851 cells after quality control. The data are normalized, scaled, and batch-aligned using preprocessing procedures implemented in the monocle3 package. Dimensionality reduction is then performed via principal component analysis (PCA), retaining 10 principal components based on the elbow plot. For external validation, each cell is annotated with one of 29 cell types and 10 tissue types, as provided in the original publication [37].

We generate three initial clustering configurations for the PHM algorithm using hierarchical clustering, DensityCut, and the Leiden algorithm, each chosen specifically for its ability to scale to the given large sample size. All methods operate on the same preprocessed data, but each produces a clustering configuration optimized under its own criterion. Notably, none of the clustering results perfectly matches the 29 annotated cell types.

The Leiden algorithm yields 17 clusters, grouping all but two of the 11 neuronal cell types into a single cluster. Similarly, pharyngeal epithelial, gland, and muscle cells are grouped together. In contrast, some cell types, such as seam cells and body wall muscle cells, are split into multiple separate clusters (Supplementary Figure 13). DensityCut also produces 17 clusters but with a different grouping structure from the Leiden algorithm. For example, all neuron types are assigned to a single cluster. Additionally, excretory, glial, and vulval precursor cells are grouped together into a cluster that also contains seam cells. Other hypodermal cells, both seam and non-seam, are distributed among multiple clusters: seam cells appear in four clusters and non-seam hypodermal cells are distributed across seven (Supplementary Figure 14). Hierarchical clustering yields only nine clusters. Neurons are split across two clusters, one predominantly composed of neurons and the other mixing the remaining neuronal cells with 11 other cell types, including excretory, coelomocyte, glial, and seam cells. As observed in DensityCut, seam and non-seam hypodermal cells are dispersed across multiple clusters (Supplementary Figure 15). Across all methods, most of the remaining annotated cell types tend to be grouped into distinct clusters. These findings suggest that when a large, diverse set of cell types is jointly analyzed, the optimal flat clustering configurations produced by different algorithms tend to be more sensitive to algorithm-specific analytical assumptions and thus less consistent.

Despite substantial differences in the initial clustering configurations, the PHM algorithm consistently reveals patterns that may reflect genuine biological relationships among the cells. Notably, the PHM results show that clusters tend to merge across cell types within the same tissue before merging across tissues (Figure 6 and Supplementary Figures 16-17), indicating a higher degree of cell type similarity within tissues. Across all initial configurations, the first merges consolidate neuron and hypodermis clusters into their respective tissue-level groups. In the Leiden-based initialization, body wall muscle cells are also merged early into a coherent group. The three prominent tissue-level clusters – neurons, hypodermis, and body wall muscle – are clearly identifiable in the PHM dendrograms and heatmaps across all cases. At intermediate resolutions, similar tissues are frequently merged in the PHM runs. For example, in both DensityCut- and Leiden-based results, the neuron group merges with the coelomocyte cluster, while the hypodermis group merges with the excretory cell clusters. A similar pattern also emerges in the hierarchical clustering initialization, although the cluster composition differs. The final merges represent the most distinct cell types, consistently involving clusters of glial cells, gonadal cells, and body wall and intestinal/rectal muscle cells. While some variation in the exact merging order exists due to differences in the initial cluster compositions, there is a strong overall consensus in the PHM-derived merging patterns across the three clustering methods.

**Figure 6:**
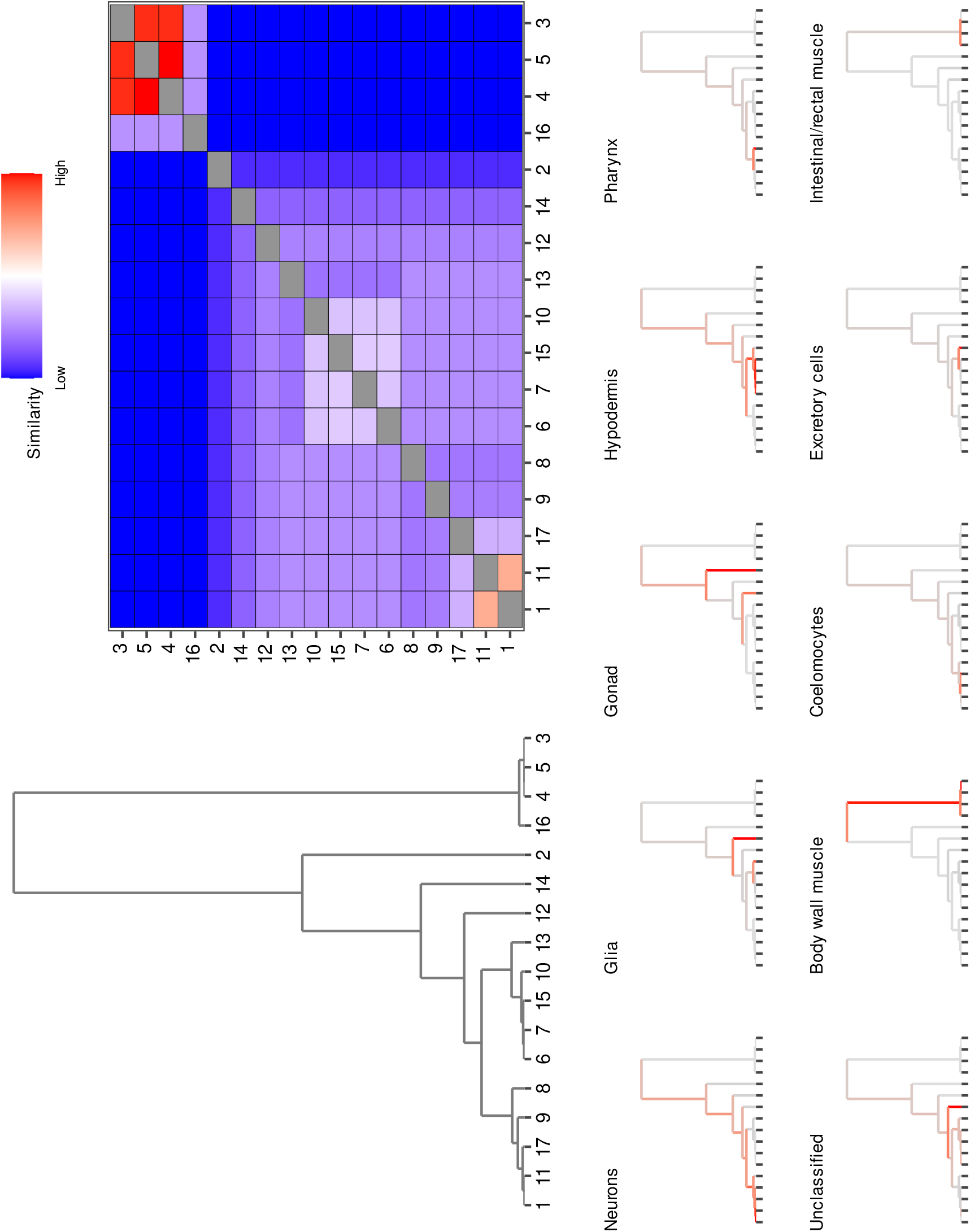
Cluster analysis of the larval *C. elegans* data. *(Top)* PHM dendrogram and heatmap from the analysis of the larval *C. elegans* cells initialized using the Leiden algorithm. *(Bottom)* The PHM dendrogram broken down by tissue types. The color indicates the proportion of cells within each branch belonging to a given tissue. Together, the dendrogram and heatmap reveal tissue-level structures emerging from the initial clustering of the data.

Although the dataset includes 11 annotated neuron subtypes, none of the clustering methods effectively recover these subtypes. We suspect this is due to limited resolution resulting from jointly preprocessing and clustering all cell types, which may obscure finer structures within specific groups. Given that neurons tend to cluster together based on the PHM results, a natural question is whether these finer substructures can be recovered by examining this subset more closely – analogous to “zooming in” on an inferred cluster. To this end, we isolate the neuron subtree identified in the PHM output based on the Leiden initialization. This subset includes 6,865 cells, of which 6,783 belong to the 11 annotated neuron subtypes and 82 to 12 non-neuronal cell types. We reapply the same preprocessing procedures as before, selecting 13 principal components based on the elbow plot. The subset is then re-clustered using the Leiden algorithm, which identifies 9 clusters. The refined clustering reveals finer structures among neuronal cell types (Supplementary Figure 18). Of the nine inferred clusters, six are predominantly composed of a single cell type, and two show near-perfect correspondence with annotated subtypes: touch receptor neurons and canal-associated neurons. Meanwhile, we observe that some annotated subtypes are split across multiple clusters, and two inferred clusters contain mixtures of six of the eleven annotated neuronal cell types, suggesting that even finer-resolution structures may still remain uncovered. Applying the PHM algorithm to quantify the relationships among the inferred neuron clusters, we find that the canal-associated neurons and three clusters enriched for ciliated sensory neurons are mostly distinguishable from other neuronal cell types at this resolution. This strategy for refining clustering structure is known as subclustering in the literature [38] and has been widely applied in scRNA-seq cluster analysis [39, 40, 41]. Here, we demonstrate that subclustering can be seamlessly integrated into our multi-resolution clustering framework with the aid of the PHM algorithm.

In summary, despite differences in initial cell-type clustering across methods, we find that the PHM procedure consistently recovers tissue-level relationships among inferred cell clusters. Moreover, the PHM-derived grouping provides a natural starting point for a divide-and-conquer, multi-stage clustering strategy: by selecting a coarse tissue-level group identified by PHM, one can repeat targeted re-clustering and the PHM procedure to uncover even finer-resolution cell-type structures.

## 3 Discussion

In this paper, we introduce a probabilistic distinguishability measure (*P*_mc_) to quantify the separability of clusters inferred from arbitrary clustering algorithms. Building on this metric, we develop the PHM algorithm to assess relative similarities among clusters and to explore induced cluster structures across multiple resolutions. When apply these methods to genetic and single-cell RNA sequencing data, and show that the PHM algorithm reveals biologically meaningful patterns that offer insights into underlying biological processes.

Examining clustering structures at multiple resolutions challenges the conventional assumption of a single optimal clustering solution, as illustrated by our “nested diamonds” example. In fact, continuous biological processes, such as human migrations and cell differentiation, naturally give rise to varying cluster patterns corresponding to different time points within each process. Rather than searching for an optimal number of clusters, the PHM algorithm focuses on sequentially merging the most similar (i.e., the closest) clusters from an initial configuration, thereby producing coarser-resolution structures. Conversely, finer-resolution structures can be identified by recursively clustering data subsets (i.e., subclustering [38]), also guided by the initial configuration and PHM results. Our findings also have implications for validating clustering results using externally annotated group labels, a common practice in benchmarking clustering methods with scRNA-seq data. As group labels may correspond to different cluster structures across resolutions, caution is warranted when interpreting instances of “over-” or “under-” clustering at any given resolution.

The PHM algorithm falls into the category of agglomerative algorithms and shares certain features with hierarchical clustering and HDBSCAN, such as the ability to produce a dendrogram. What fundamentally distinguishes these algorithms is the choice of similarity measures used to guide cluster merging, which leads to substantial differences in outcomes, as demonstrated in our double ring and real data examples. The probabilistic similarity measure implemented by *P*_mc_ offers some unique conceptual and practical advantages, facilitating more intuitive interpretation of PHM results. Importantly, its broad applicability allows the PHM algorithm to operate on the output of *any* clustering method, providing a principled and independent means to quantify relative relationships among inferred clusters.

It is well recognized that clustering results generated by different algorithms can appear inconsistent, as demonstrated in our synthetic and real data examples. In addition to the fact that different algorithms often operate at different resolutions, such discrepancies also stem from the lack of a precise and universally accepted definition of a “cluster.” Clustering algorithms are typically guided by distinct criteria, such as centroid proximity, density, or connectivity, which reflect different conceptualizations of what constitutes a cluster and may not align in all scenarios. The PHM algorithm has demonstrated the ability to identify reproducible patterns across outputs from diverse clustering methods. Such concordant patterns, emerging from methods based on fundamentally different model assumptions, are arguably more robust and more likely to capture underlying biological truths. This property is particularly valuable in unsupervised learning, where external validation sources can be limited. Building on this strength, the PHM algorithm can be further developed to support fully automated *ensemble clustering*, which naturally extends our current work.

Finally, we acknowledge that cluster analysis of genomic data – particularly scRNA-seq data – is inherently complex. While the broad compatibility and adaptability of the PHM algorithm make it straightforward to integrate into existing single-cell analysis pipelines, the effects of upstream preprocessing steps, such as normalization and dimension reduction, on the outcomes of multi-resolution clustering remain insufficiently understood. In our future work, we will investigate these aspects more systematically, with the goal of contributing to the development of best practices for single-cell data analysis.

## 4 Methods

### 4.1 Distinguishability Criterion *P*_mc_

The Distinguishability criterion, *P*_mc_, measures the overall separability among all clusters in a given configuration. It is defined by quantifying the probability of misclassifying a data point away from its generating cluster in a multi-class classification problem. Below, we describe the setup and derivation of *P*_mc_ in detail and discuss its key properties, which motivate the development of the PHM algorithm.

#### 4.1.1 Defining *P*_mc_

Given a configuration of *K* clusters, we denote the generating class label of a data point ***x*** by *θ ∈* 1, …, *K*. We assume a pre-defined classifier *δ*(***x***) : ℝ^*p*^ *→* 1, …, *K* that assigns ***x*** to its *predicted* generating class. The performance of the classifier can be evaluated using the 0-1 loss function, defined as

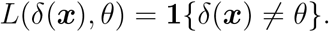

We use *P*_mc_ to quantify the average misclassification probability across all possible data points and generating clusters in the given configuration. Formally, *P*_mc_ is defined as the Bayes risk [42] of the classifier *δ*(***x***) in the statistical decision theory, i.e.,

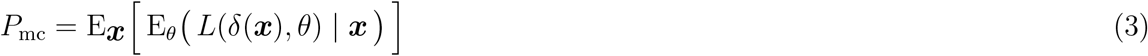

The use of 0-1 loss ensures that the resulting Bayes risk is a valid probability measure ranging from 0 to 1. It can be naturally interpreted as the probability of misclassifying a data point under the given clustering configuration, averaged over all possible values of ***x*** and their true generating clusters.

As our use of *P*_mc_ is solely for comparing the separability of different clustering configurations, any valid classifier *δ*(***x***) can be applied within this framework. For improved interpretability and computational convenience, our implementation focuses on classifiers that operate directly on the probabilities

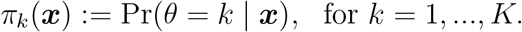

This set includes the optimal classifier under the 0-1 loss, *δ*_*o*_, defined as,

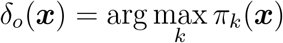

Our default classifier for computing *P*_mc_ is a randomized decision function, *δ*_*r*_, which assigns a label to an observation ***x*** by sampling from a categorical distribution based on the probability vector ***π***(***x***) = (*π*_1_(***x***), …, *π*_*K*_(***x***)), i.e.,

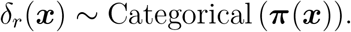

In addition to producing *P*_mc_ values that closely align with those from the optimal classifier in decision-relevant ranges of cluster separation (Supplementary Figure 19), the randomized classifier’s computational properties ensure the high efficiency of the PHM algorithm.

For a given data point ***x***, *π*_*k*_(***x***) is evaluated using the Bayes rule,

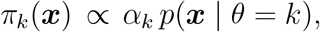

where *α*_*k*_ denotes the prior probability of cluster *k* and *p*(***x*** | *θ* = *k*) is the likelihood function.

The likelihood function encodes the characteristics of the corresponding cluster population and is evaluated using the cluster-specific density estimation result detailed in Section 4.1.3. The prior quantifies the relative frequency of the observations arising from each assumed cluster. By default, we estimate the relevant priors from the observed clustering data, but they can also be set to user-specified values if desired.

#### 4.1.2 Computation of *P*_mc_

The Bayes risk for a general classifier *δ*(***x***) : ℝ^*p*^ *→* {1, …, *K*} assigning a point ***x*** to one of the *K* clusters can be derived as follows,

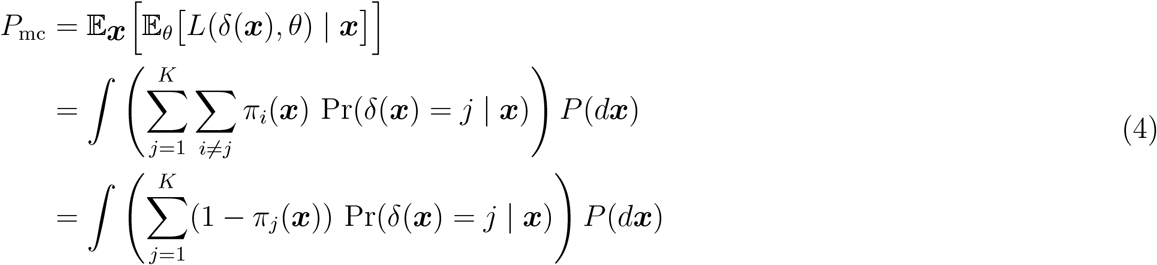

For a more detailed derivation, see Supplementary Methods 2. The marginal data distribution *P* (***x***) is given by

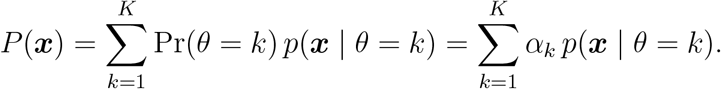

For the default randomized decision rule, *δ*_*r*_(***x***) *∼* Categorical (***π***(***x***)),

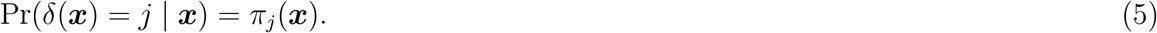

Thus

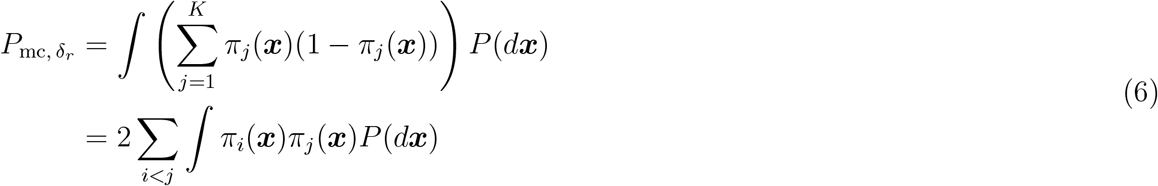

To compute *P*_mc_ using the optimal decision rule, *δ*_*o*_(***x***) = arg max_*k*_ *π*_*k*_(***x***), we define a partition of the sample space, 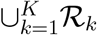, such that ℛ_*k*_ := {***x*** : *δ*_*o*_(***x***) = *k*}. It follows that,

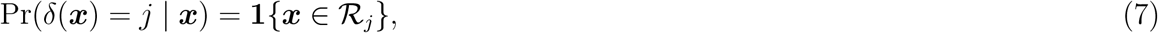

and

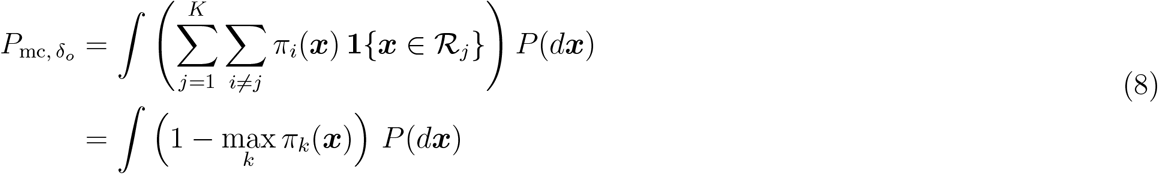

Note that,

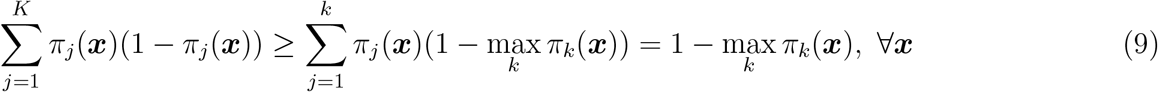

Hence,

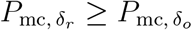

Unless otherwise specified, we use the notation *P*_mc_ to refer to 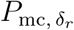 by default.

#### 4.1.3 Density Estimation for Partitioned Clustering Data

Computing *P*_mc_ requires an estimate of the underlying cluster densities. Unlike model-based clustering approaches that produce these densities as a byproduct, many commonly used hard clustering methods only yield a partition of the observations. A straightforward but naïve strategy is to estimate the cluster-specific densities (e.g., using a GMM) based solely on the subset of observations assigned to each cluster. However, this approach ignores uncertainty in the cluster assignments and can result in an underestimation of *P*_mc_.

To address this problem, we propose the following procedure for estimating the density of each partition. Specifically, we model the distribution of the cluster population corresponding to each partition as a mixture of Gaussian densities. To account for the uncertainty in hard clustering assignment, the procedure begins by computing the cluster membership probability for each observation. We first define the distance from an observation *x*_*i*_ to a partition *C*_*k*_ as the minimum distance between *x*_*i*_ and any observation in *C*_*k*_:

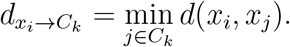

If *x*_*i*_ is originally assigned to partition *C*_*k*_, this distance is 0; otherwise, 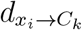 is strictly greater than 0. Based on this distance metric, we compute the weight of *x*_*i*_ representing *C*_*k*_ using a softmax function:

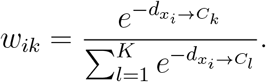

There are alternative options for defining the point-to-partition distance 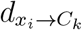 and *w*_*ik*_. The aforementioned naïve approach can be viewed as defining *w*_*ik*_ = 0 if *x*_*i*_ is assigned to *C*_*k*_, and 1 otherwise. Another viable option is to use the average distance between *x*_*i*_ and all points in partition *C*_*k*_. Finally, the resulting weights are incorporated into a weighted EM algorithm [43] to estimate the densities associated with each partition.

A comparison the naïve and weighted cluster density estimates for the double ring example is shown in Supplementary Figure 20.

#### 4.1.4 Lower and Upper Bounds of *P*_mc_

When all clusters are well-separated, *P*_mc_ approaches its lower bound of 0. More specifically, in the case of well-separated clusters, the following two conditions hold:

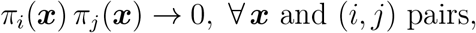

and

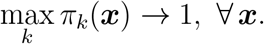

Hence, both decision rules, *δ*_*r*_ and *δ*_*o*_, yield perfect classification accuracy, which can be straight-forwardly derived from Eqns (6) and (8), respectively.

At the other extreme, *P*_mc_ is maximized when all clusters are completely overlapping, i.e.,

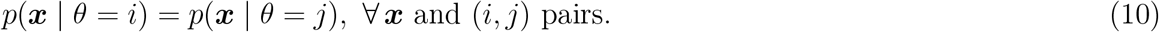

Thus,

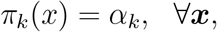

and

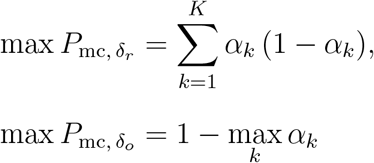

In the special case of equal priors (i.e., *α*_*k*_ = 1*/K* for all *k* values), it follows that

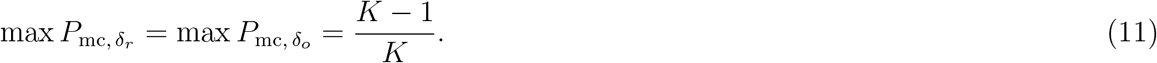

#### 4.1.5 The Cluster Merging Property of *P*_mc_

The merging property is specific to the default *P*_mc_, which is evaluated using the randomized decision rule *δ*_*r*_. For a given clustering configuration with *K ≥* 2, consider merging two arbitrary clusters to form a new, combined cluster. Let *P*_mc_ and 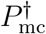 denote the misclassification probabilities before and after the merging, respectively. The following proposition summarizes this merging property concerning the reduction in the overall misclassification probability.

##### Proposition 1.

*Merging two existing clusters indexed by i and j leads to*

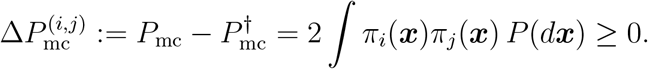

*Furthermore*,

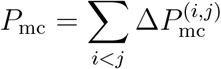

*Proof*. Supplementary Methods 2 □

The cluster merging property forms the basis of the PHM algorithm. Since 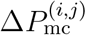 can be pre-computed for all pairs of clusters from the initial configuration, subsequent updates to *P*_mc_, caused by merging one pair of clusters at a time, become straightforward to compute.

#### 4.1.6 Numerical Evaluation of *P*_mc_

Numerical evaluation of *P*_mc_ can be challenging, especially when the clustering data are high-dimensional. For low-dimensional data, it is possible to evaluate Eqn. (4) using numerical integration with various quadrature methods. However, these methods generally do not scale well when the data dimensionality exceeds 5. When the marginal data distribution *P* (***x***) can be sampled from directly (as is the case in all examples presented in this paper), Monte Carlo (MC) integration provides an efficient alternative. Specifically, we draw *M* data points from *P* (***x***) and approximate *P*_mc_ by

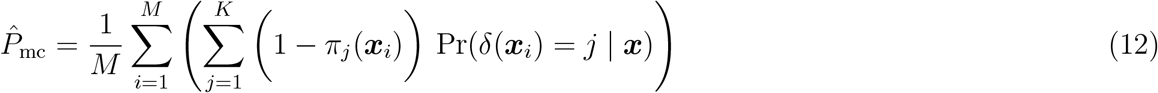

One of the key advantages of the Monte Carlo method is that its error bound remains 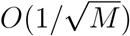 regardless of the dimensionality of ***x***. We provide comparisons between the Monte Carlo and numerical integration methods for evaluating *P*_mc_ in selected low-dimensional settings (Supplementary Methods 3), demonstrating that MC integration is both accurate and efficient.

### 4.2 The PHM algorithm and Visualizations

#### 4.2.1 The PHM algorithm

The *P*_mc_ hierarchical merging (PHM) algorithm constructs a hierarchy of clusters based on an initial clustering configuration. The similarity between clusters–which guides the merging process–is characterized by their pairwise Δ*P*_mc_ values. Large values of Δ*P*_mc_ indicate low distinguishability and a high degree of overlapping between clusters. The algorithm proceeds by iteratively merging the least distinguishable clusters, i.e., those with the largest 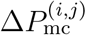, until only a single cluster remains. Since 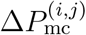 can be precomputed for all cluster pairs from the initial configuration, subsequent updates for *P*_mc_ are straightforward by using the cluster merging property of *P*_mc_ (Proposition 1). The complete algorithm, including Δ*P*_mc_ update steps, is shown in Algorithm 1.

##### Algorithm 1: *P*_mc_ Hierarchical Merging (PHM) Algorithm

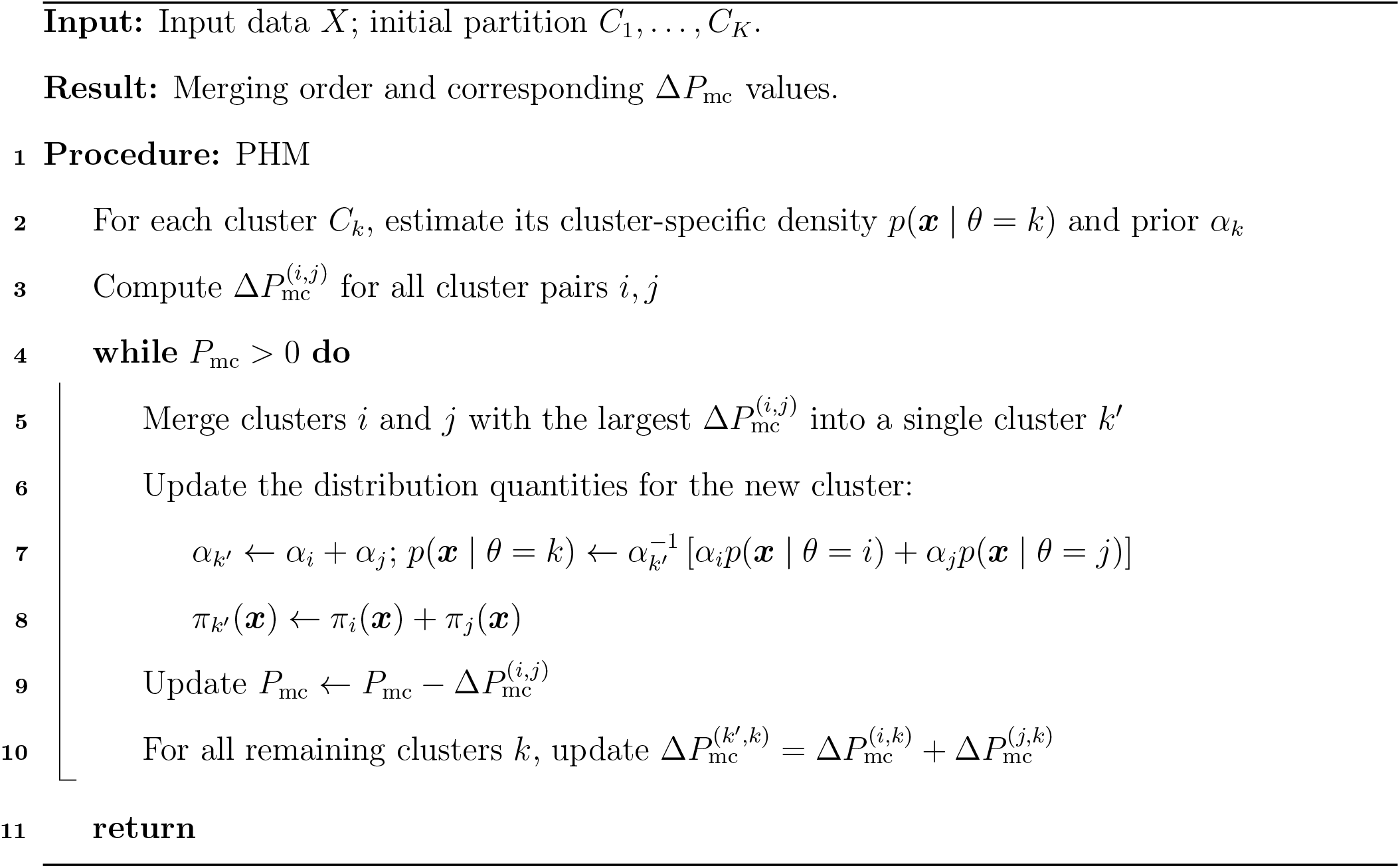

#### 4.2.2 Dendrogram Visualization

Given the hierarchical nature of the merging process, it is natural to visualize the PHM algorithm using a dendrogram. Consider a subtree 𝒮 with *m* leaf nodes (original clusters) formed by merging subtrees 𝒮_1_ and 𝒮_2_. By default, the height of the merge for subtree 𝒮 is set inversely proportional to:

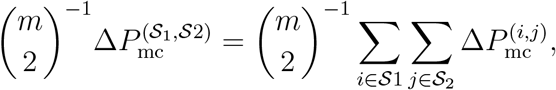

which reflects the average dissimilarity among the original clusters being merged.

Alternatively, the merge height can be computed based on the maximum dissimilarity between any pair of clusters in the subtrees. In this case, we set the branch height inversely proportional to:

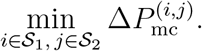

For visualization purposes, a log 10 transformation of the branch heights is applied by default to better reveal different clustering resolutions.

#### 4.2.3 Heatmap visualization

For initial configurations including many clusters, it may be clearer to visualize the PHM algorithm using a heatmap that captures the relationships between all pairs of initial clusters. Specifically, each heatmap cell corresponding to the original clusters *i* and *j* is based on 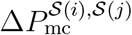, which represents the Δ*P*_mc_ value at which these clusters are first combined into the same merged cluster. When multiple resolutions are present in the data, these appear as visible block-diagonal patterns in the heatmap, as shown in Sections 2.2.1 and 2.4.

By default, we apply a log 10 transformation to construct the heatmap color scale, i.e., the heatmap is based on:

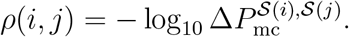

Alternatively, because log_10_ Δ*P*_mc_ is not always linear with respect to distance (Supplementary Figure 19), we also provide an option to use a spline-based transformation to scale the Δ*P*_mc_ values so they roughly correspond to the distance between two univariate Gaussian clusters (Supplementary Methods 4) for the color scale.

### 4.3 Clustering Methods

Here we briefly summarize the clustering methods used to generate the initial clustering configurations for the PHM procedure.

#### 4.3.1 *k*-means clustering

Clusters in *k*-means clustering are characterized by their centroids, the mean of all observations in the cluster. The clustering is usually performed using Lloyd’s algorithm [44], which alternates assigning observations to the cluster with the nearest centroid and updating the cluster centroid to eventually arrive at a locally optimal solution for a specified number of clusters *K*. The “optimal” value of *K* is chosen using the Silhouette statistic, an external validation criterion which compares the within-cluster distances to the between cluster distances, to determine the “best” clustering. We use the *k*-means and silhouette statistic [45] implementations from the `ClusterR` [46] R package.

#### 4.3.2 Hierarchical clustering

Hierarchical clustering produces a sequence of partitions for observations, starting from the partition where each observation is its own cluster. Hierarchical clustering uses a pre-specified linkage function (common examples include single linkage, complete linkage, and the Ward linkage) to define the distance between clusters, and at each step merges the clusters with the smallest distance between them. This hierarchy, visualized using a dendrogram, can be “cut” to obtain a partition for a given number of clusters *K*. The “optimal” value of *K* is chosen using the Silhouette statistic, an external validation criterion which compares the within-cluster distances to the between cluster distances, to determine the “best” clustering. We use the hierarchical clustering implementation from the `fastcluster` [47] R package.

#### 4.3.3 Finite mixture models

Finite mixture models (FMM) use a mixture of distributions, with a finite number of components *k* specified in advance, to fit the overall distribution of the data. [48, 49] Once fit to the data, each component of the FMM is taken as its own cluster. However, rather than produce a partition of the data, the FMM produces a posterior probability distribution over the cluster assignments for each observation. A post-hoc classification procedure, with a pre-specified decision rule using the cluster assignment probabilities, can be applied to partition the observed data.

The Gaussian mixture model (GMM) is probably the most commonly used mixture model in practice because of its flexibility to fit many diverse data types [29], and is our representative FMM approach in our analysis. We use the Bayesian information criterion (BIC) to determine the number of components to fit the data. We use the GMM implementation from the Mclust [43] R package.

#### 4.3.4 Density-based Clustering

Density-based clustering relies on the intuition that clusters correspond to “high density” regions of the data, in which many observations are in close proximity to one another, and are separated by “low density” regions that are sparse in the data. Because of this, density-based approaches do not need to have the number of clusters specified beforehand, and are generally robust to different cluster shapes. The representative approach for density-based clustering we adopt is DenstiyCut [50], which uses a refined *k*-nearest neighbor density estimate to identify. DensityCut is an improved density-based clustering algorithm derived from the widely used DBSCAN/OPTICS algorithm [51], which is also included in our synthetic data analysis. DensityCut compares the density differences at and between modes of the estimated distribution to obtain a cluster stability measure, which it uses to determine its cluster structure. We use the DensityCut implementation from the `DensityCut` [52] R package.

#### 4.3.5 Community Detection

Community detection is commonly used to find clusters within graphs, which are characterized by a high degree of within-group connection and a low degree of out-of-group connection [23, 22]. The Leiden and Louvain algorithms construct clusters agglomeratively based on a modularity criterion, which compares the degree of within-cluster connectivity to the between cluster connectivity, combining clusters which increase the modularity until no further improvements can be made. Our representative approaches for community detection are the Leiden and Louvain algorithms, for which we use the implementations from the `monocle3` [53] and `Seurat` [54] packages respectively.

### 4.4 Real Data Analysis Details

#### 4.4.1 Human Genome Diversity Project

For preprocessing, we first construct an allele dosage matrix for the individuals comprising this data. Next, we perform principal component analysis on the allele dosage matrix. For each of the first 100 and 200 PCs (which capture 70% and 85% of the variability in the data, respectively), we determine the elbow point using the `fildElbowPoint` function from the `PCAtools` [55] R package. This dimension reduced data is used as input to the clustering methods.

We fit the varying clustering methods to the data as follows. For the GMM we compare the BIC of models fit with 1-20 mixture components and all supported covariance structures (unstructured, diagonal, equal variance, etc) implemented in the `Mclust` package. For *k*-means and hierarchical clustering, we choose *K* based on the clustering with the best average silhouette scores [45], implemented in the `clusterR` package, searching between 1-20 clusters. We run the Leiden and DensityCut clustering methods using their default settings. Lastly, when running the HDBSCAN procedure we use a minPts value of 5, which is the suggested default value.

#### 4.4.2 Peripheral Blood Mononuclear Cells

Data are preprocessed using the standard workflow implemented in the Seurat package. We first discard cells with very few (<200) or a high number (>2500) of expressed genes, as well as cells with a high percentage (>5%) of reads mapping to mitochondrial genomes. We then perform normalization, scaling, and PCA dimension reduction using the following Seurat functions: `NormalizeData`, `FindVariableFeatures`, `ScaleData`, and `RunPCA`. We determine the elbow point from the PCA embedding based on the first 200 PCs using the `fildElbowPoint` function from the `PCAtools` R package. The PCA embedding is then extracted and used as input to the clustering methods under consideration.

We fit the varying clustering methods to the data as follows. For the GMM we determine the best model by comparing the BIC of models fit with 1-15 mixture components and all supported covariance structures implemented in the Mclust package. For *k*-means and hierarchical clustering, we choose the value of *K* corresponding to the clustering with the largest average silhouette score, searching from between 1-20 clusters. The DensityCut and Leiden procedures are run using the default settings, and for HDBSCAN we use a value of 5 for the minPts parameter.

#### 4.4.3 Larval *C. elegans* Cell Profiling

Data are processed using the standard workflow implemented in the monocle3 package. Specifically, the data normalization, scaling, PCA dimension reduction, and batch alignment steps are performed using the `preprocess_cds` and `alig_cds` functions. We determine the elbow point based on the first 100 PCs using the `fildElbowPoint` function from the `PCAtools` R package. The PCA embedding is then extracted and used as input to the clustering methods under consideration. For the follow-up analysis of the neuron subtree, we repeat these steps on that subset of the data.

We fit the varying clustering methods to the data as follows. We use the default settings for the Leiden and DensityCut clustering procedures. For hierarchical clustering, we choose the value of *K* based on the average silhouette score, searching from between 1-60 clusters.

