## Supplementary Material for "Multiresolution Clustering of Genomic Data"

### Supplementary Figures

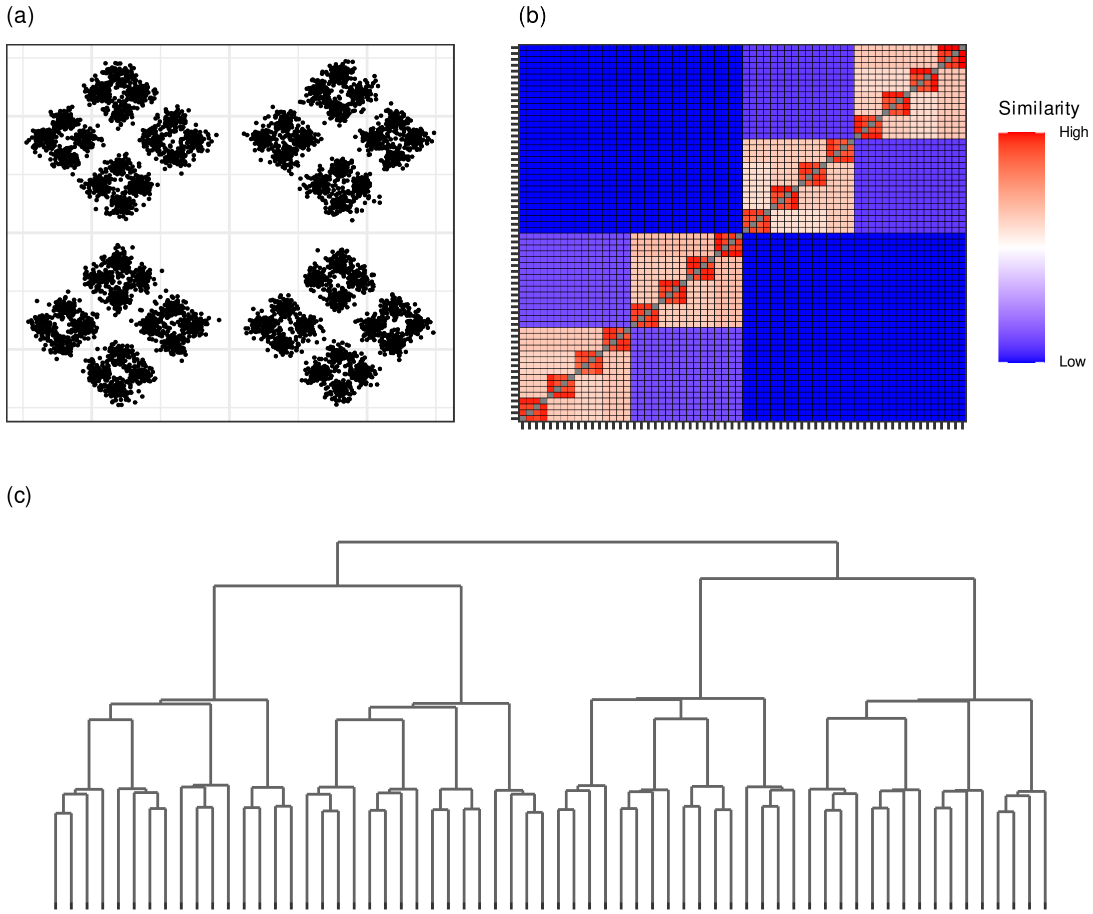


Supplementary Figure 1: **PHM detects multi-level clustering structures in the nested diamonds (DensityCut)** (a) Simulated data from 64 clusters arranged in a nested diamonds pattern. Clusters are arranged in three layers of structure: inner diamonds (4 original clusters), outer diamonds (4 inner diamonds), and a single outer square. (b) Heatmap visualizing the values of $\Delta P_{\mathrm{mc}}$ at the branches in the PHM dendrogram initialized from the DensityCut solution reveals three distinct levels of cluster structure. (c) The dendrogram visualizing the PHM procedure, with subtree heights grouping at three levels, corresponding to the three levels of structure in this data.

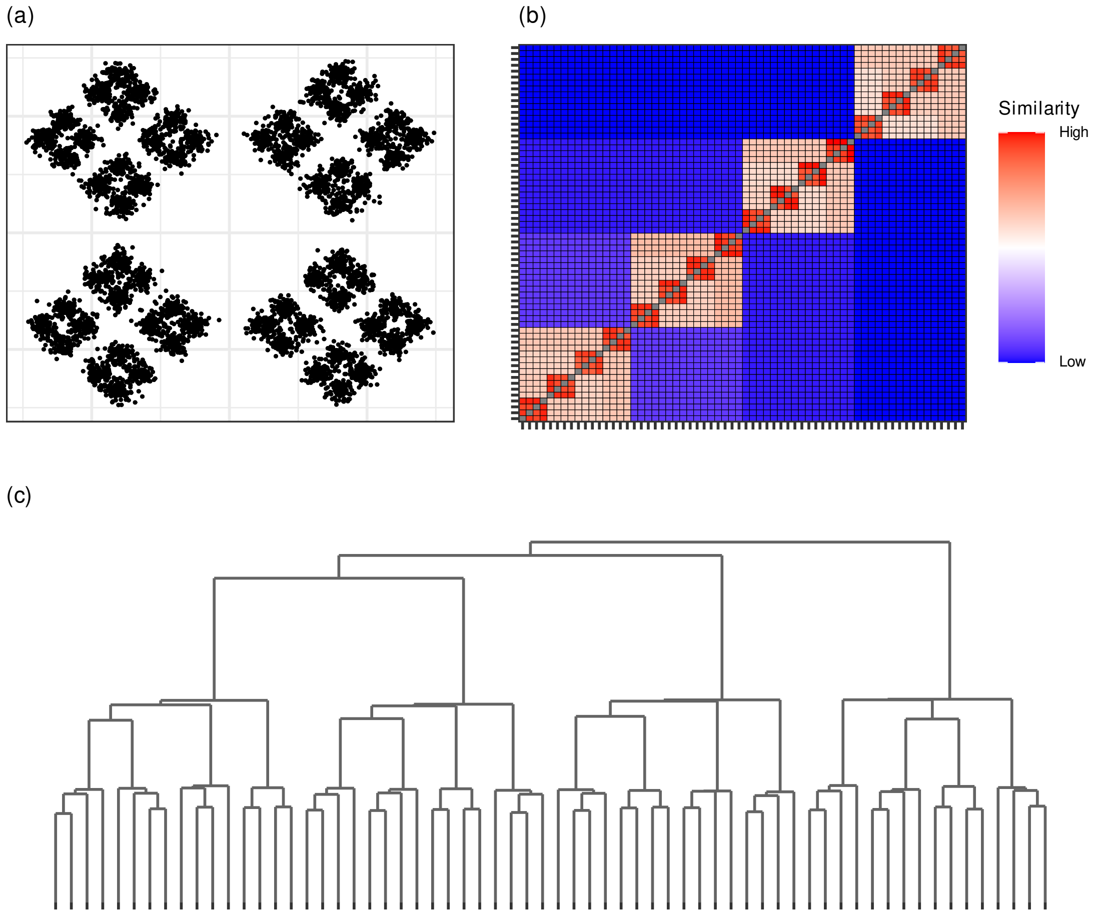


Supplementary Figure 2: **PHM detects multi-level clustering structures in the nested diamonds (*k*-means)** (a) Simulated data from 64 clusters arranged in a nested diamonds pattern. Clusters are arranged in three layers of structure: inner diamonds (4 original clusters), outer diamonds (4 inner diamonds), and a single outer square. (b) Heatmap visualizing the values of $\Delta P_{\mathrm{mc}}$ at the branches in the PHM dendrogram initialized from the $k$-means solution with $K = 64$ reveals three distinct levels of cluster structure. (c) The dendrogram visualizing the PHM procedure, with subtree heights grouping at three levels, corresponding to the three levels of structure in this data.

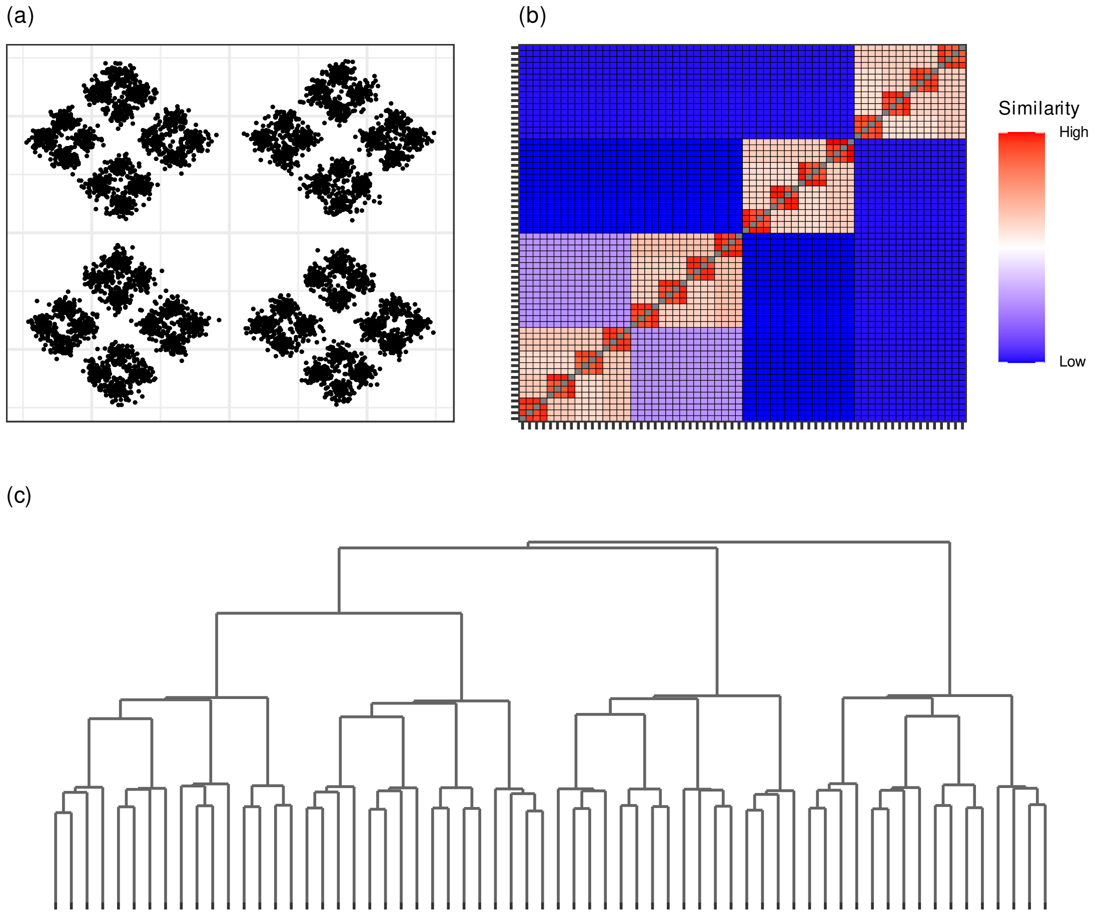


Supplementary Figure 3: **PHM detects multi-level clustering structures in the nested diamonds (hierarchical clustering)** (a) Simulated data from 64 clusters arranged in a nested diamonds pattern. Clusters are arranged in three layers of structure: inner diamonds (4 original clusters), outer diamonds (4 inner diamonds), and a single outer square. (b) Heatmap visualizing the values of $\Delta P_{\text{mc}}$ at the branches in the PHM dendrogram initialized from the hierarchical clustering solution with $K = 64$ reveals three distinct levels of cluster structure. (c) The dendrogram visualizing the PHM procedure, with subtree heights grouping at three levels, corresponding to the three levels of structure in this data.

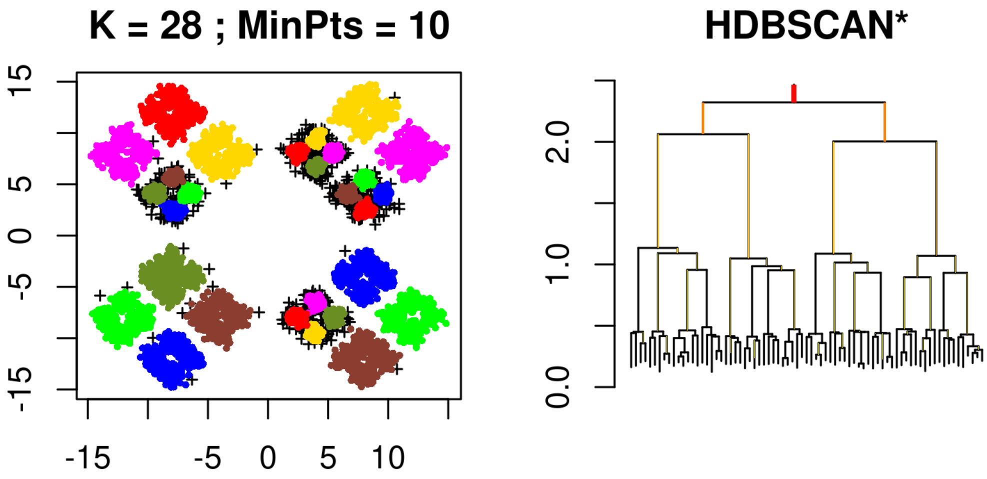


Supplementary Figure 4: **HDBSCAN recovers multi-level structure in nested diamonds** *(Left)* The chosen partition from the HDBSCAN algorithm using $minPts = 10$, which tends to capture the structure in the data at the resolution of the "inner diamonds." Some of the original clusters are over-partitioned. *(Right)* Resulting dendrogram from the HDBSCAN algorithm, from which the three layers of structure in the data can be seen based on the clustering of the merge heights into three distinct groups.

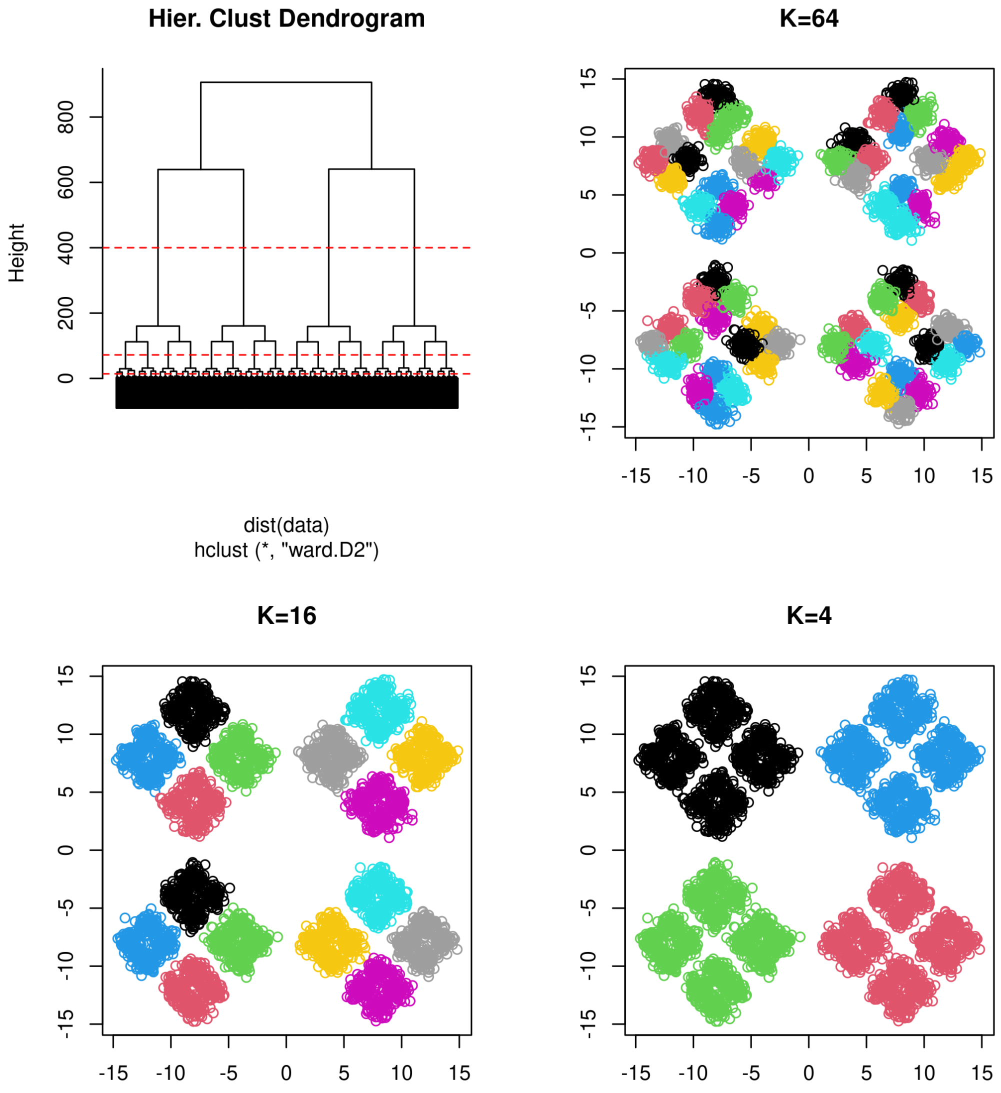


Supplementary Figure 5: **Hierarchical clustering recovers multi-level structure in nested diamonds** *(Top Left)* Dashed red lines on the dendrogram correspond to cuts at $K = 4, 16, 64$. *(Top Right)* Cluster assignments when $K = 64$ roughly captures the highest-resolution grouping of the observations into individual clusters. *(Bottom Left)* Cluster assignments when $K = 16$ captures the grouping of clusters into the 16 "inner diamonds." *(Bottom Right)* Cluster assignments when $K = 64$ captures the grouping of clusters into the 4 "outer diamonds."

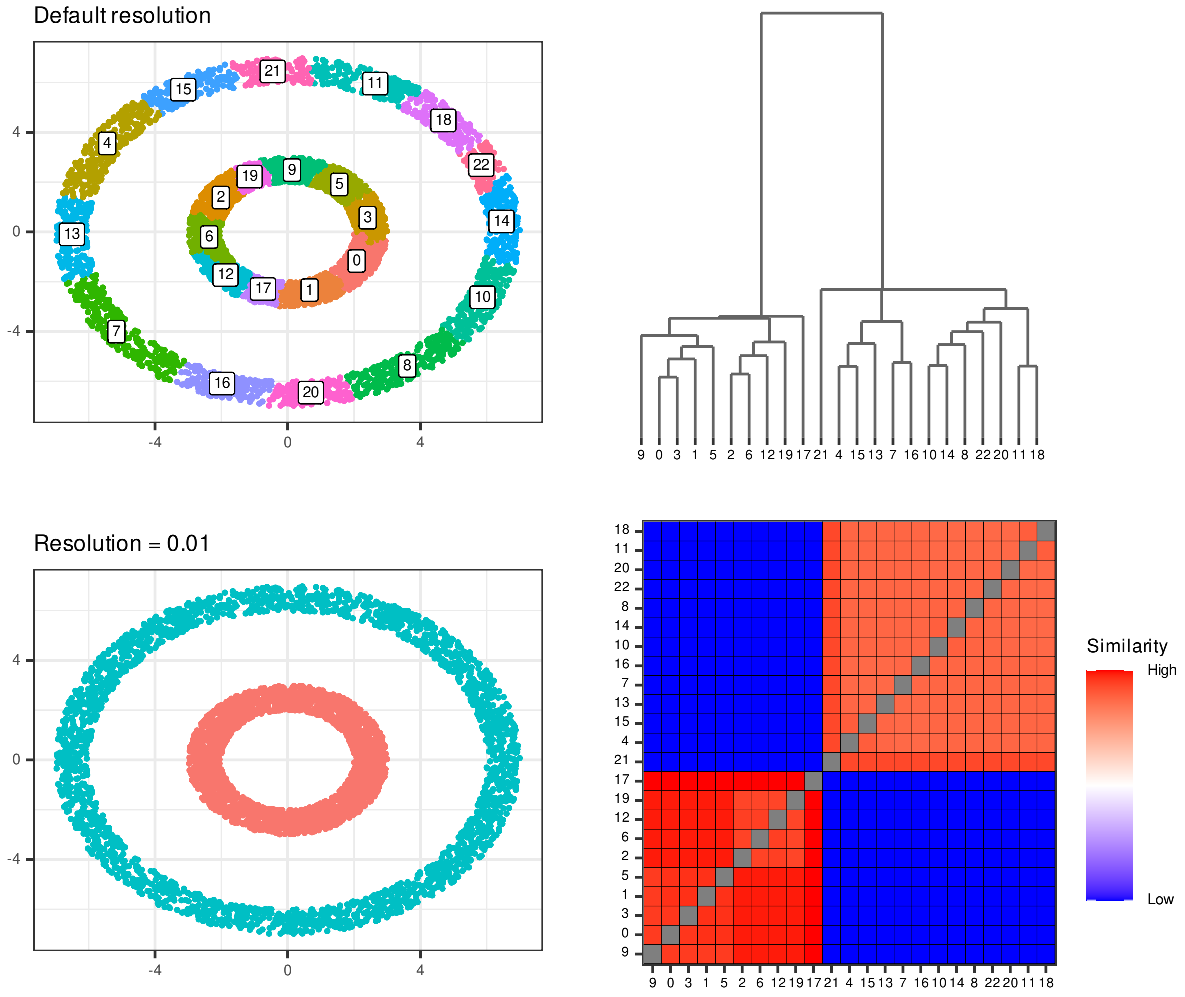


Supplementary Figure 6: **PHM recovers the double-ring structure from Louvain clustering** *(Top Left)* Louvain clustering results under default parameters for the double ring simulated example. The Louvain algorithm highly over-clusters this data, identifying 23 communities. *(Bottom Left)* Louvain clustering results with resolution parameter = 0.01 identifies two distinct communities, corresponding to each of the rings. *(Top Right)* PHM demdrogram based on the default Louvain clustering results. *(Bottom Right)* PHM heatmap visualization based on the default Louvain clustering results. The PHM algorithm aggregates the clusters into two distinct structures, corresponding to each of the rings.

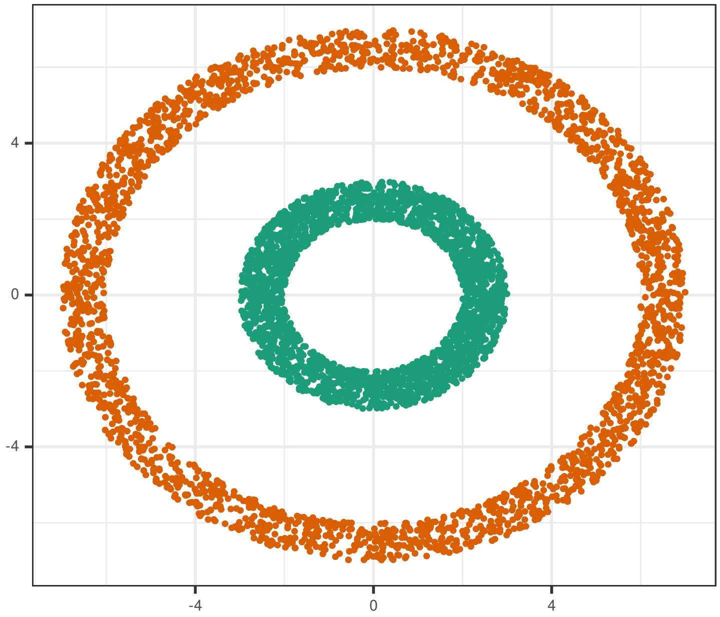


Supplementary Figure 7: **DensityCut results for the double ring example** DensityCut clustering results for the double ring simulated data example. DensityCut, with default parameters, is correctly able to identify the two rings as distinct clusters in this example.

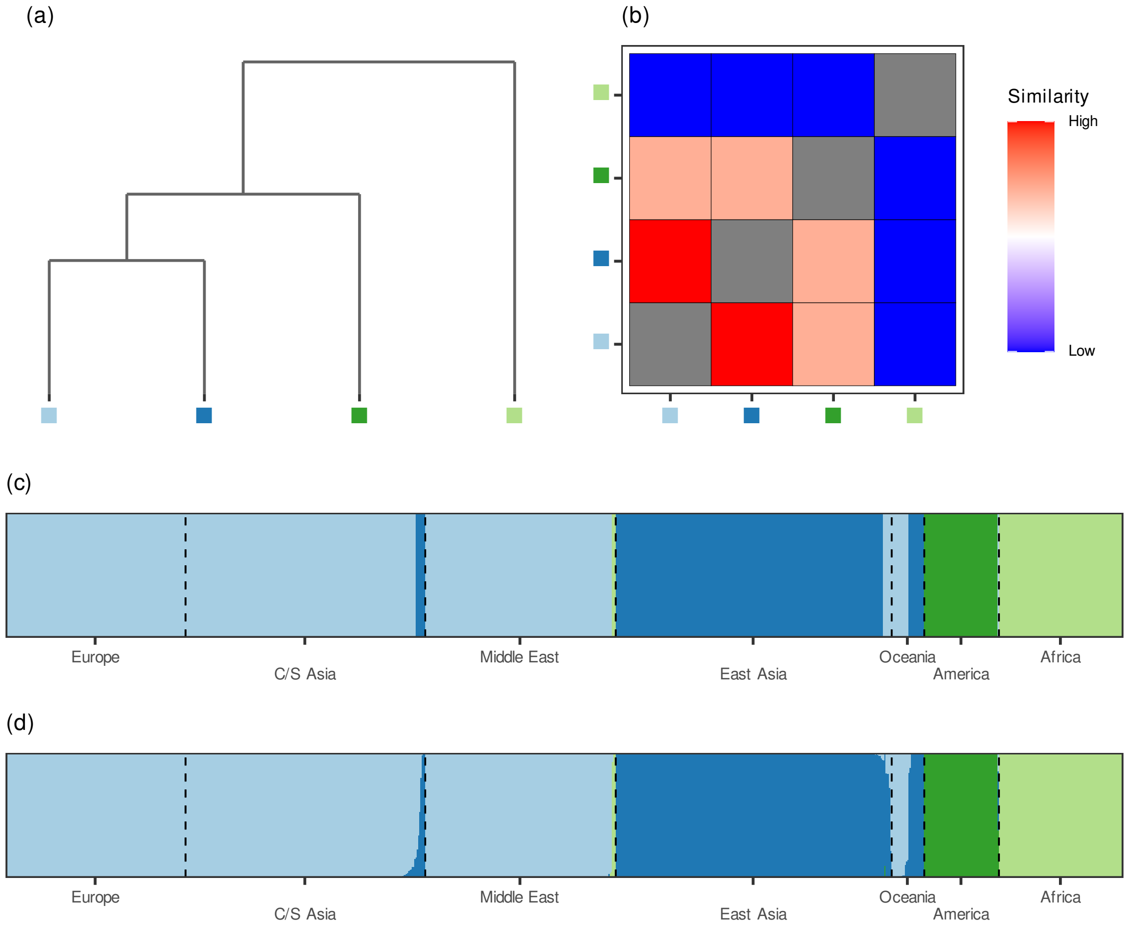


Supplementary Figure 8: **PHM result from HGDP data analysis (DensityCut)** (a) PHM dendrogram visualizing the merging of the 4 clusters estimated from DensityCut. (b) PHM heatmap visualizing the $\Delta P_{\rm mc}$ values from the merging procedure. (c) Distruct plot [1] visualizing the cluster assignments for each observation grouped by geographic sampling location. (d) Posterior probabilities for each observation based on the estimated cluster-specific densities.

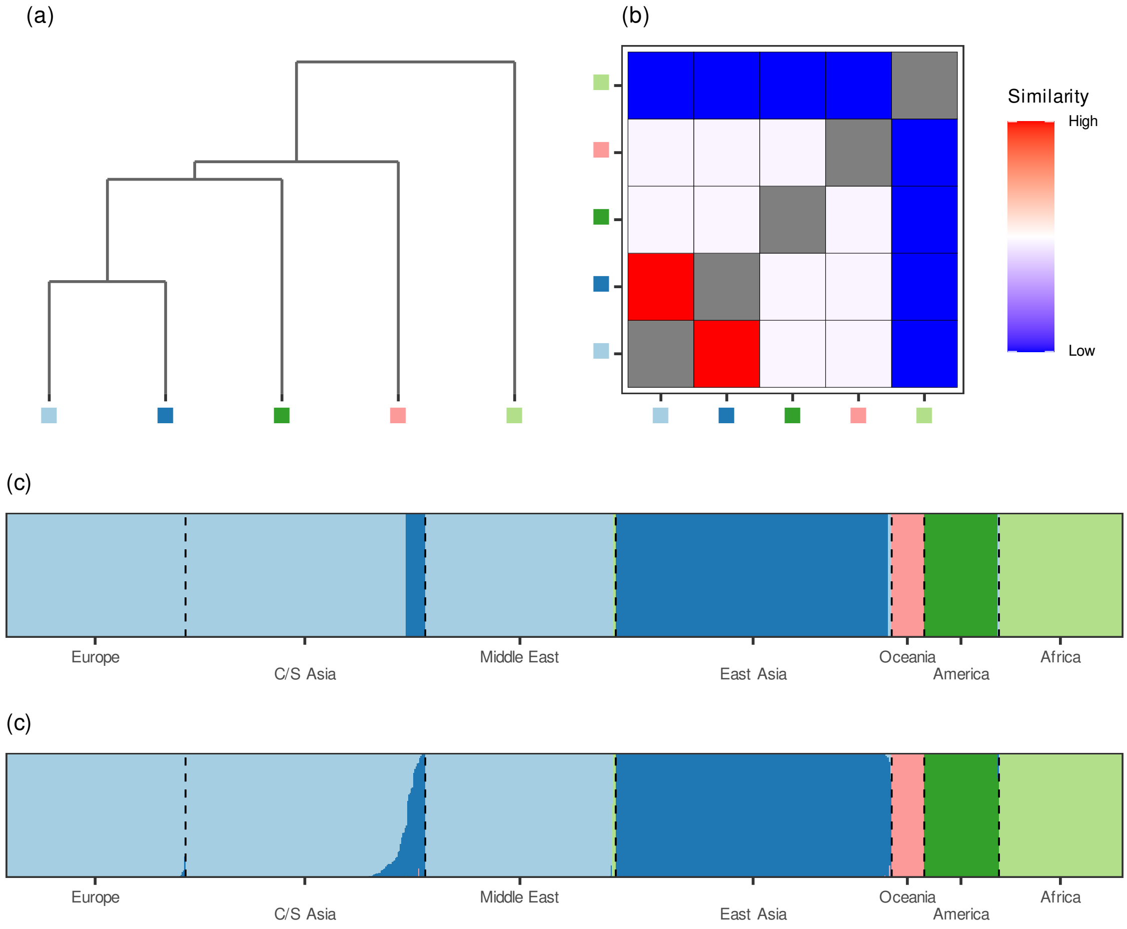


Supplementary Figure 9: **PHM result from HGDP data analysis (hierarchical clustering)** (a) PHM dendrogram visualizing the merging of the 5 clusters estimated from hierarchical clustering. (b) PHM heatmap visualizing the $\Delta P_{\mathrm{mc}}$ values from the merging procedure. (c) Distruct plot [1] visualizing the cluster assignments for each observation grouped by geographic sampling location. (d) Posterior probabilities for each observation based on the estimated cluster-specific densities.

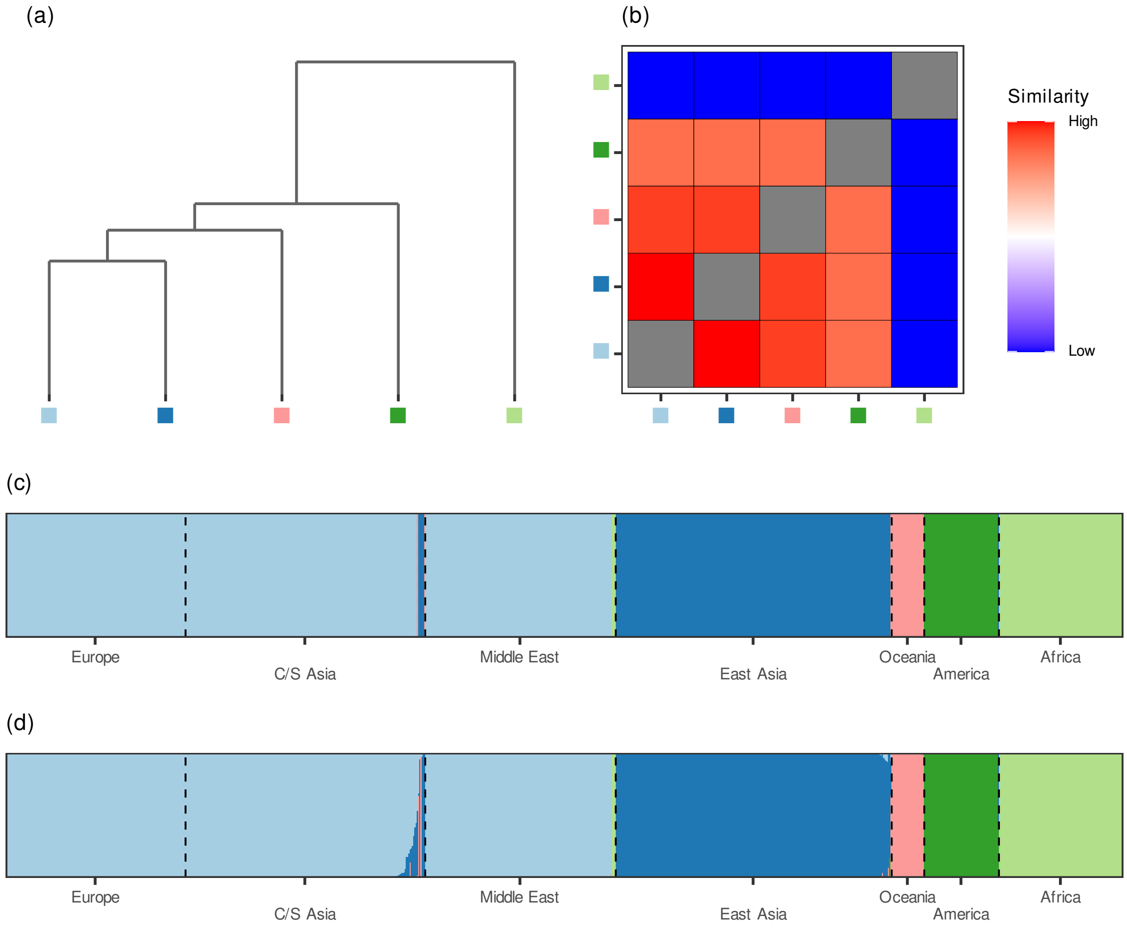


Supplementary Figure 10: **PHM result from HGDP data analysis (*k*-means)** (a) PHM dendrogram visualizing the merging of the 5 clusters estimated from *k*-means. (b) PHM heatmap visualizing the $\Delta P_{\text{mc}}$ values from the merging procedure. (c) Distruct plot [1] visualizing the cluster assignments for each observation grouped by geographic sampling location. (d) Posterior probabilities for each observation based on the estimated cluster-specific densities.

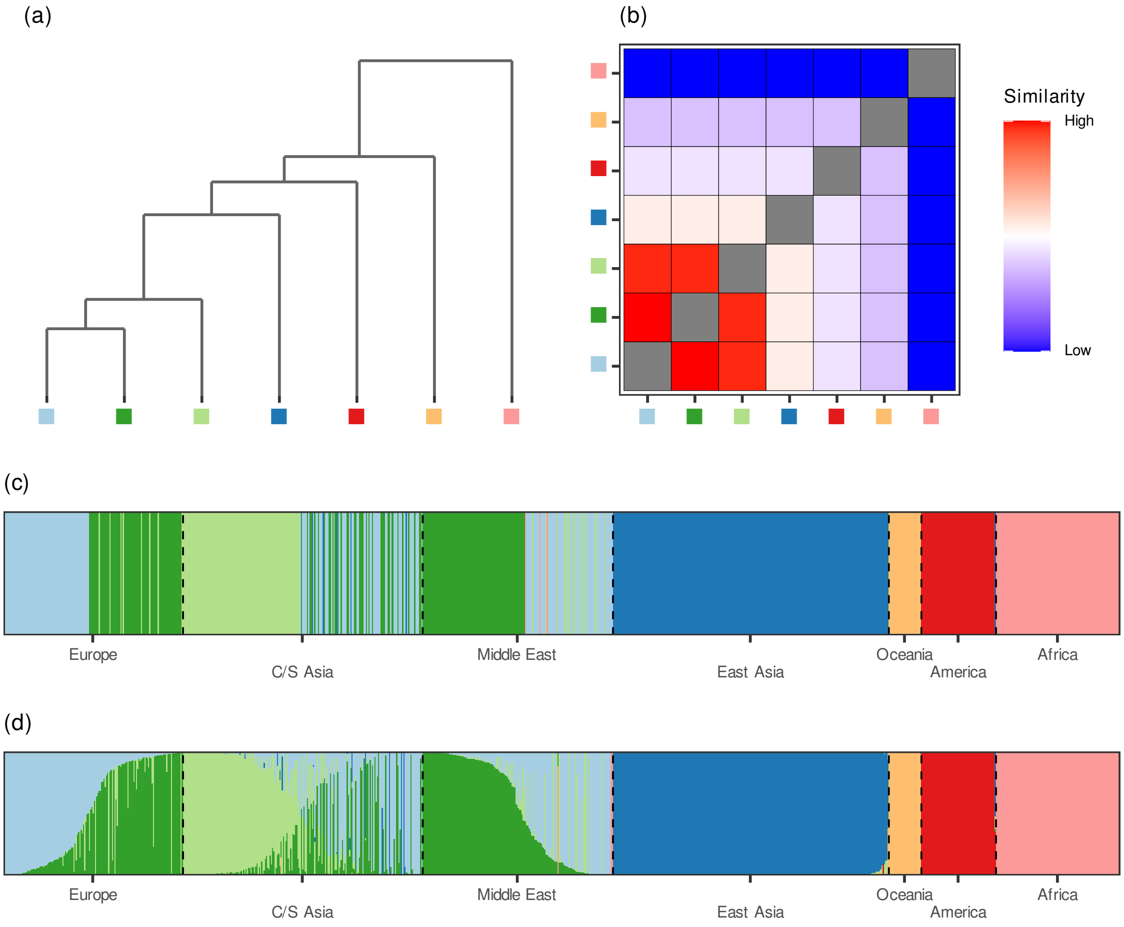


Supplementary Figure 11: **PHM result from HGDP data analysis (Leiden)** (a) PHM dendrogram visualizing the merging of the 7 clusters estimated from the Leiden algorithm. (b) PHM heatmap visualizing the $\Delta P_{\text{mc}}$ values from the merging procedure. (c) Distruct plot [1] visualizing the cluster assignments for each observation grouped by geographic sampling location. (d) Posterior probabilities for each observation based on the estimated cluster-specific densities.

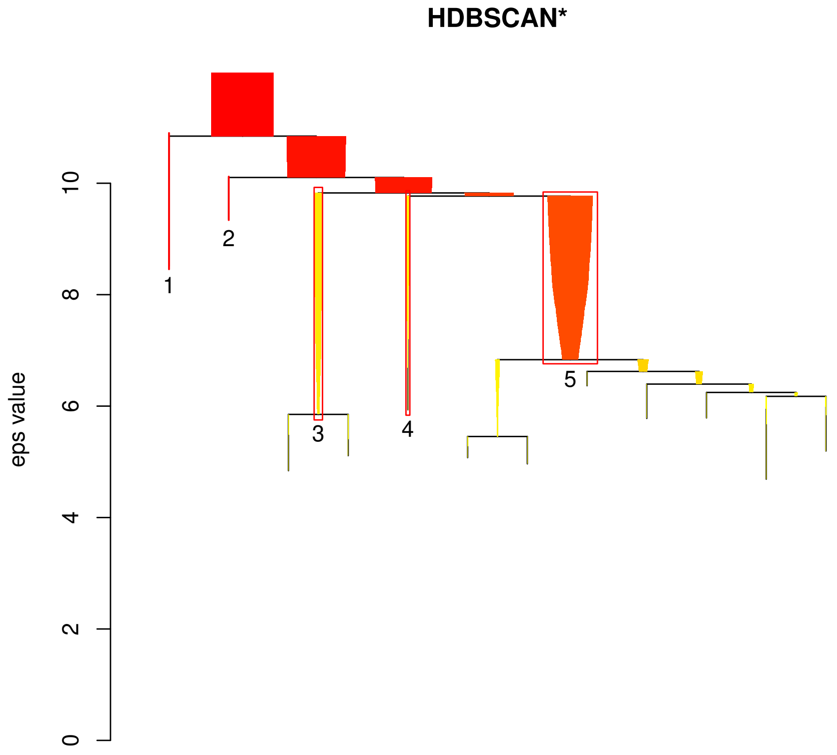


Supplementary Figure 12: **HDBSCAN result from HGDP data analysis** Cluster dendrogram from the HDBSCAN clustering procedure on the HGDP data. Boxed branches correspond to the final inferred clusters from the clustering procedure.

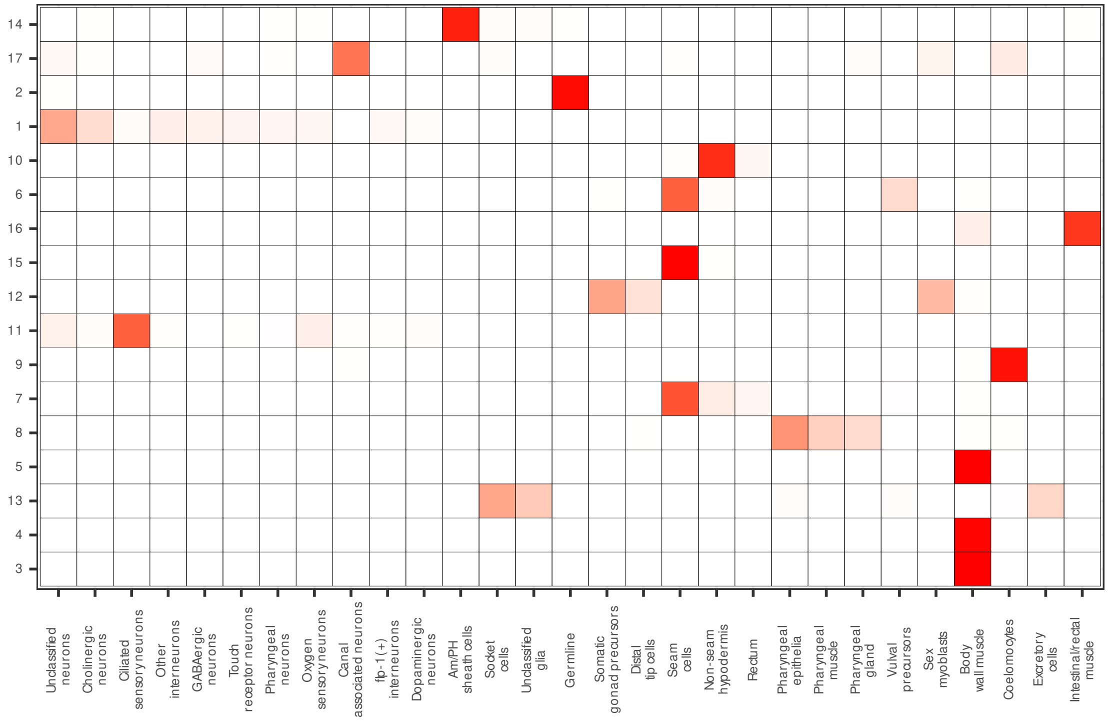


Supplementary Figure 13: **Leiden algorithm cluster composition for *C. elegans*** Cell-type distribution for the larval *C. elegans* data across clusters inferred from the Leiden algorithm. Color indicates the proportion of cells within a cluster belonging to a given cell type, ranging from red (large proportion of cluster belongs to given cell type) to white (no member of the cluster belongs to the given cell type).

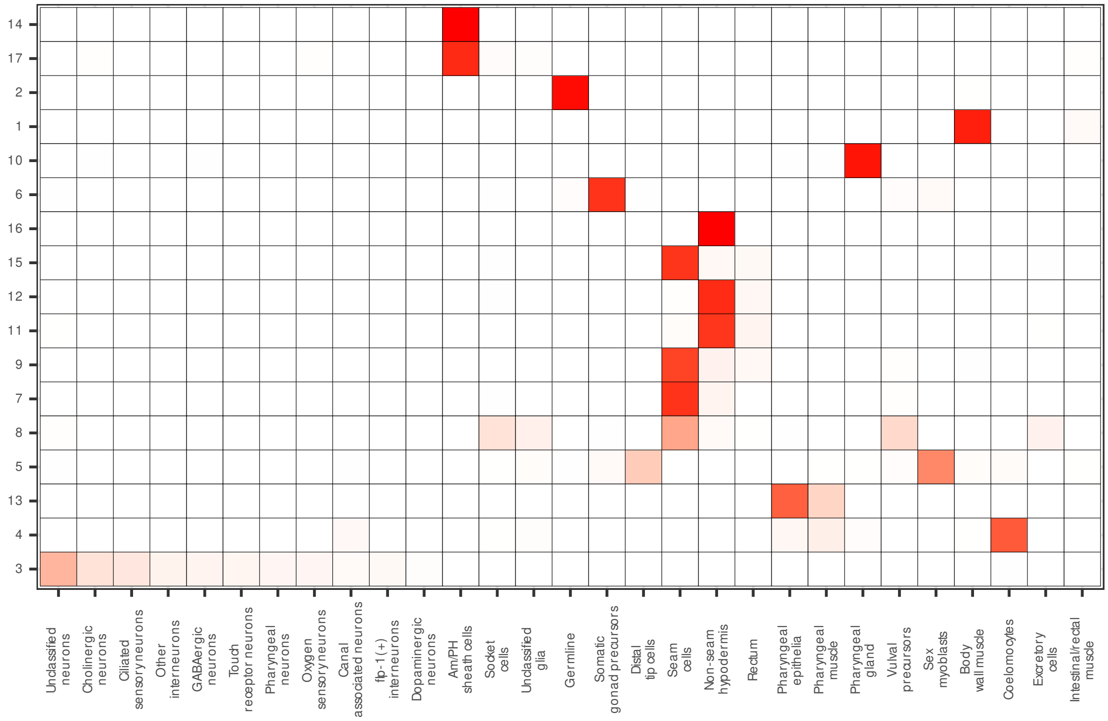


Supplementary Figure 14: **DensityCut cluster composition for *C. elegans*** Cell-type distribution for the larval *C. elegans* data across clusters inferred from DensityCut. Color indicates the proportion of cells within a cluster belonging to a given cell type, ranging from red (large proportion of cluster belongs to given cell type) to white (no member of the cluster belongs to the given cell type).

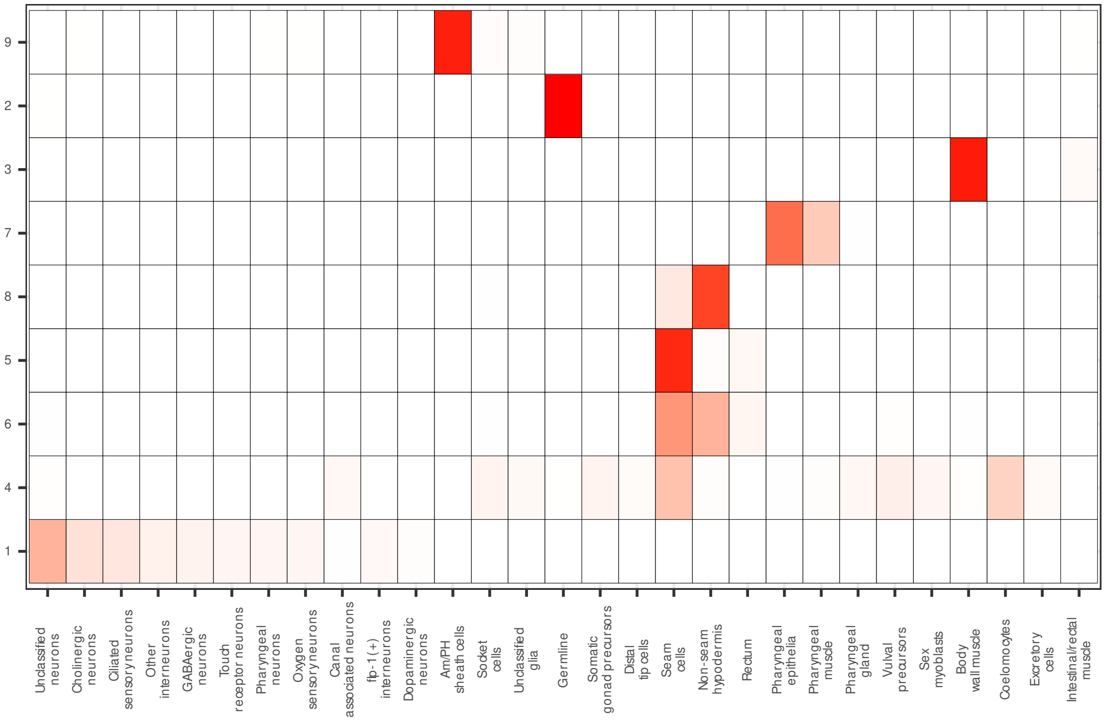


Supplementary Figure 15: **Hierarchical clustering cluster composition for *C. elegans*** Cell-type distribution for the larval *C. elegans* data across clusters inferred from hierarchical clustering. Color indicates the proportion of cells within a cluster belonging to a given cell type, ranging from red (large proportion of cluster belongs to given cell type) to white (no member of the cluster belongs to the given cell type).

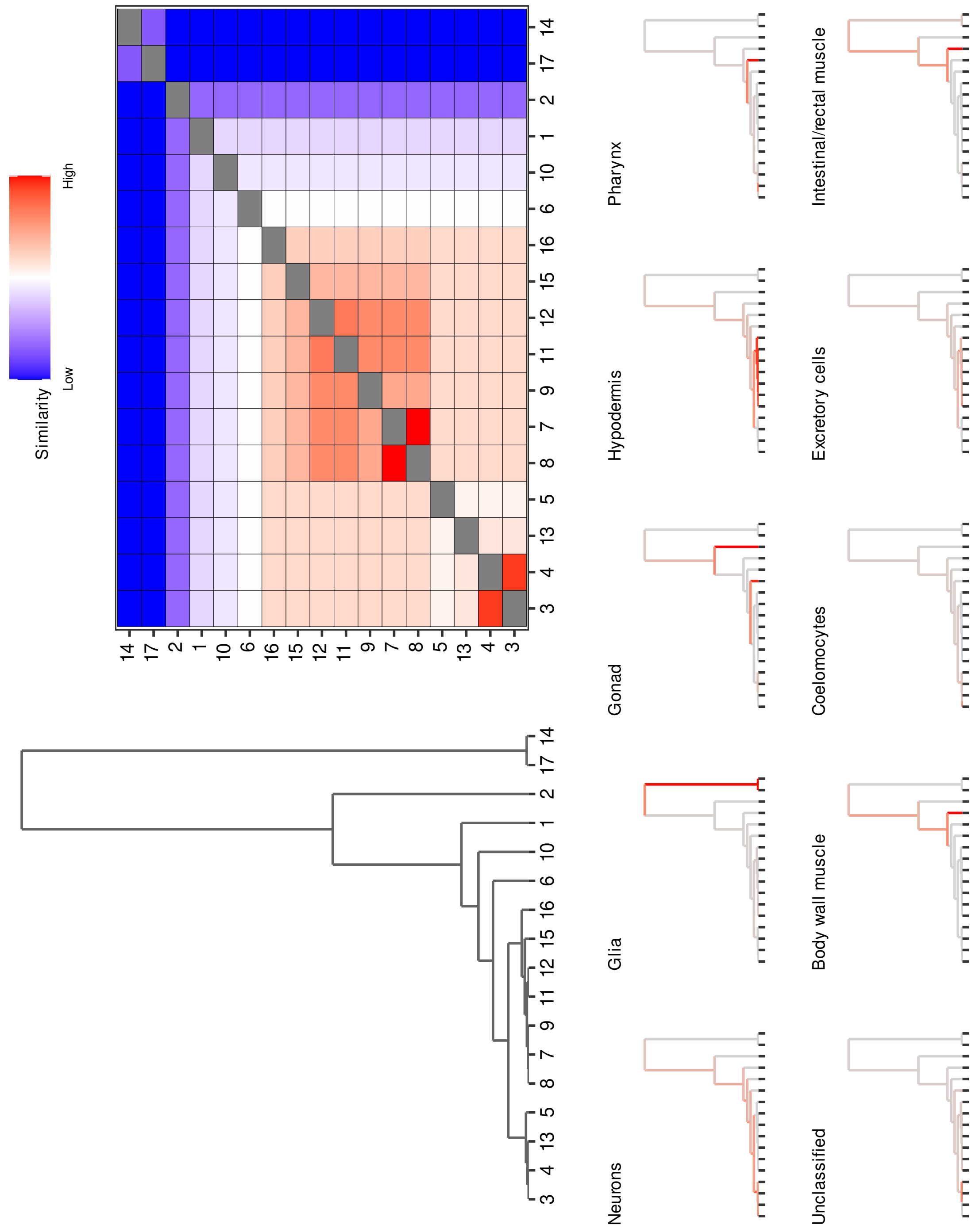


Supplementary Figure 16: **Cluster analysis of the larval *C. elegans* data (DensityCut)** *(Top)* PHM dendrogram and heatmap for the analysis of the larval *C. elegans* cells based on DensityCut clustering. *(Bottom)* The PHM dendrogram broken down by tissue types. The color corresponds to the proportion of cells in a given branch belonging to a given tissue type. The dendrogram and heatmap reveal tissue-level structures based on the initial clustering of the data.

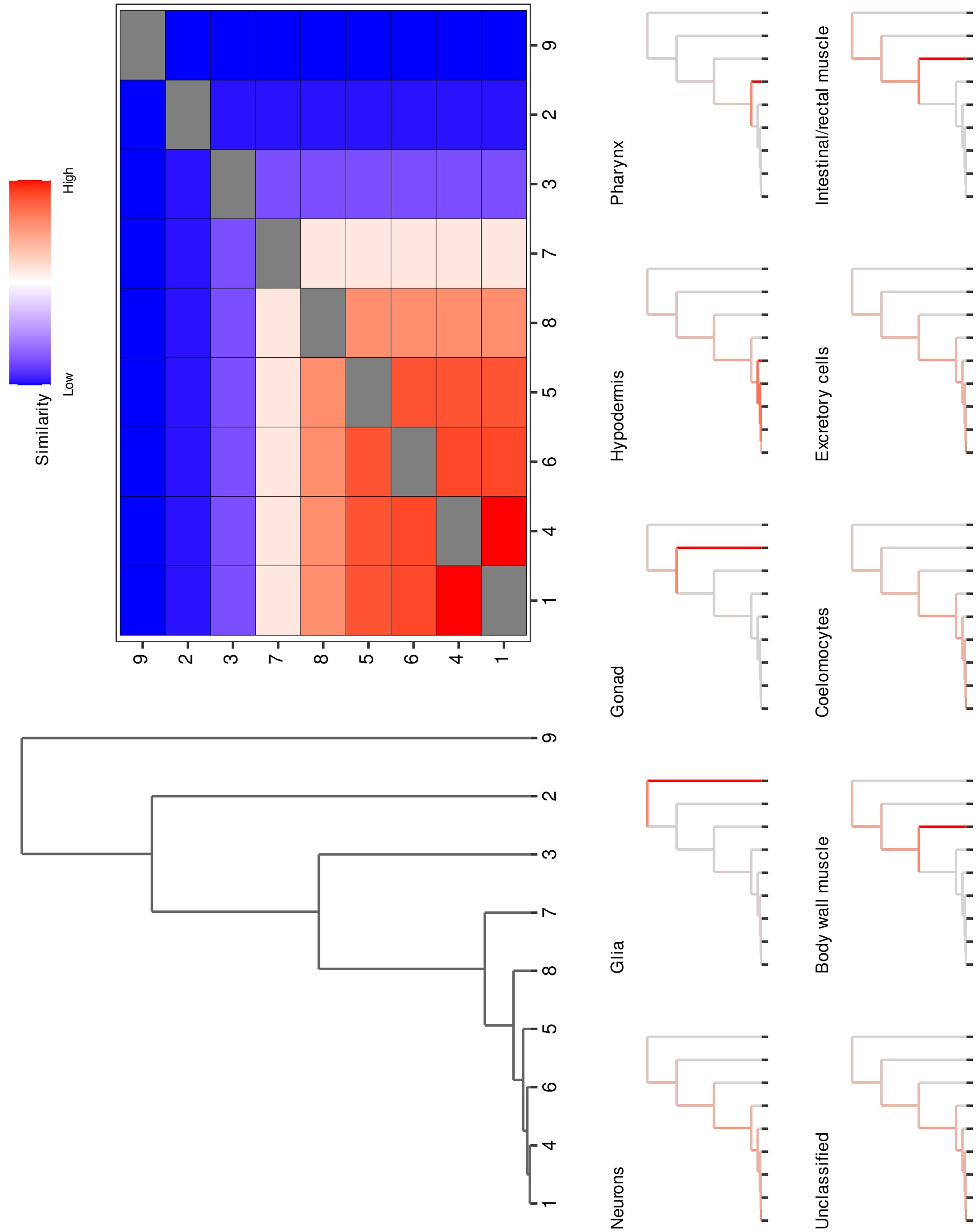


Supplementary Figure 17: **Cluster analysis of the larval *C. elegans* data (hierarchical clustering)** *(Top)* PHM dendrogram and heatmap for the analysis of the larval *C. elegans* cells based on hierarchical clustering. *(Bottom)* The PHM dendrogram broken down by tissue types. The color corresponds to the proportion of cells in a given branch belonging to a given tissue type. The dendrogram and heatmap reveal tissue-level structures based on the initial clustering of the data.

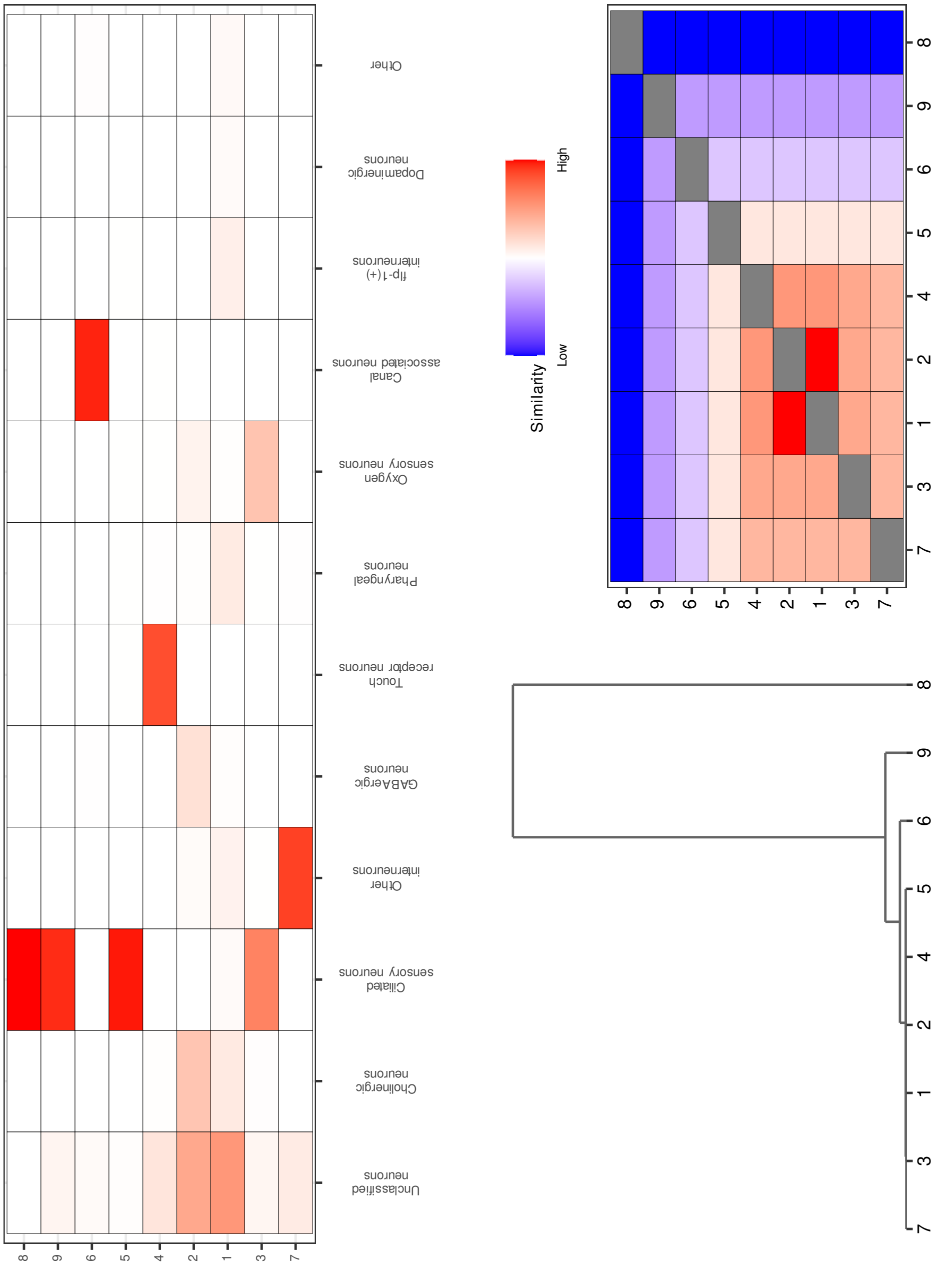


Supplementary Figure 18: **Cluster analysis and PHM results for the *C. elegans* neuron drill-down** *(Top)* Cell-type distribution across clusters from the neuron-specific drill-down of the *C. elegans* data. Color indicates the proportion of cells within a cluster belonging to a given cell type, ranging from red (large proportion of cluster belongs to given cell type) to white (no member of the cluster belongs to the given cell type). *(Bottom)* PHM dendrogram and heatmap for the neuron-specific drill-down.

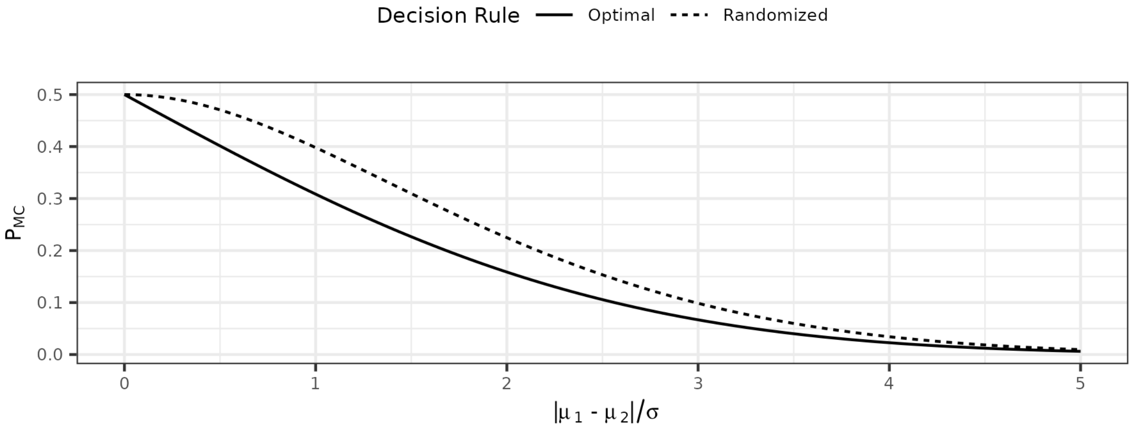


Supplementary Figure 19: $P_{\mathrm{mc}}$ **values across decision rules** Values of Pmc based on the randomized and optimal decision rules $\delta_r$ and $\delta_o$. The value $P_{\mathrm{mc}}$ is shown in the y-axis and is calculated for two univariate Gaussian distributions $N(\mu_1, \sigma)$ and $N(\mu_2, \sigma)$ where $\pi_1 = \pi_2 = 0.5$. The x-axis indicates the degree of cluster separation in terms of the distribution parameters.

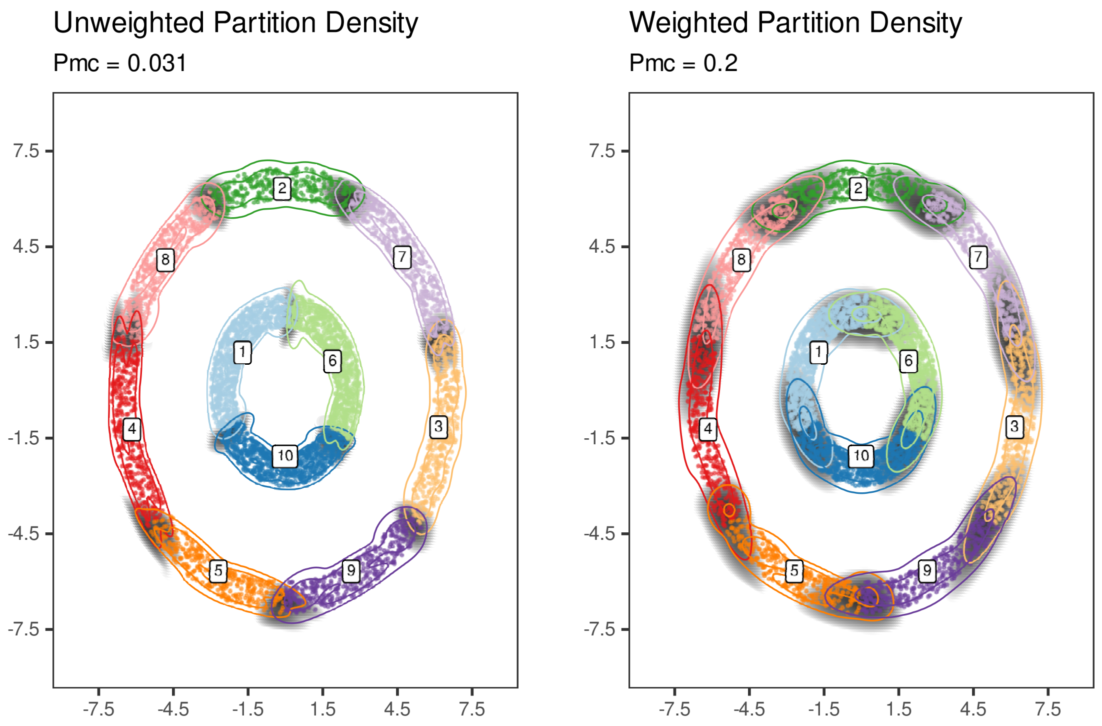


Supplementary Figure 20: **Comparison of naïve and weighted density estimation for double rings** Countour plots for the estimated densities using the *(Left)* naïve and *(Right)* weighted density estimation procedures for the $k$-means partition of the double ring simulated data example. Shaded regions indicate areas in the density contributing to $P_{\mathrm{mc}}$.

### Supplementary Tables

| | Europe | C/S Asia | Middle East | East Asia | Oceania | America | Africa |
|---|---|---|---|---|---|---|---|
| *Noise* | 7 | 21 | 16 | 16 | 14 | 12 | 0 |
| 1 | 0 | 0 | 0 | 0 | 7 | 0 | 0 |
| 2 | 0 | 0 | 0 | 0 | 6 | 0 | 0 |
| 3 | 0 | 0 | 2 | 0 | 0 | 0 | 103 |
| 4 | 0 | 0 | 0 | 0 | 0 | 49 | 0 |
| 5 | 142 | 178 | 140 | 213 | 0 | 1 | 0 |

Supplementary Table 1: HGDP observations broken down by assigned HDBSCAN cluster. The *Noise* cluster are observations not assigned to any cluster by the HDBSCAN procedure.

### Supplementary Methods

### 1 Simulation Details

#### 1.1 Nested Diamonds

To illustrate the PHM algorithm's ability to capture cluster structures at multiple data resolutions, we simulate data in a nested diamonds structure as described in Section 2.2.1. Specifically, 100 observations are drawn from 64 2-D Gaussian clusters with $0.5 \times I_2$ covariance and cluster mean $\mu_k$ determined in the following manner based on the index $k$

1. The position along the corners of the outer square $\mu_k^{(1)}$ is set to one of $\mu_k^{(1)} = \{\pm 8, \pm 8\}$
2. The position along the corners of the outer diamond $\mu_k^{(2)}$ is set to one of $\mu_k^{(2)} = \{0, \pm 4\}$ or $\{\pm 4, 0\}$
3. The position along the corners of the inner diamond $\mu_k^{(3)}$ is set to one of $\mu_k^{(3)} = \{0, \pm 1.5\}$ or $\{\pm 1.5, 0\}$
4. The cluster center can be found to be $\mu_k = \mu_k^{(1)} + \mu_k^{(2)} + \mu_k^{(3)}$, of which there are 64 possible values

The $k$-means and GMM clustering solutions can be sensitive to the initial state of the cluster parameters. For illustrative purposes, the $k$-means solution is initialized using the known cluster centers, and then fit using the standard procedure. For the GMM we initialize the procedure using the model-based hierarchical clustering approach implemented in the `Mclust` package, specifying a spherical covariance structure for the model components to ensure that the underlying cluster structure is appropriately captured.

#### 1.2 Double Ring

To illustrate the PHM algorithm's ability to identify consensus structures from seemingly disparate clustering solutions, we simulate data in a double ring structure as described in Section 2.2.2. We draw 6,000 observations in total from two rings of radius 3 (for the inner ring) and 7 (for the outer ring), where each observation has associated radius $r_i$ (either 3 or 7) and angle $\theta_i$. Each observation's radius then has a random value $u_i$ subtrated from it, where $u_i \sim Unif(0, 1)$.

GMM, $k$-means, and hierarchical clustering all search between 2 and 10 clusters for their solutions. The GMM clustering solution is chosen based on the BIC, while $k$-means and hierarchical clustering are chosen based on the averaged silhouette score. The Louvain and DensityCut results are performed using the default parameters, however we use a resolution parameter of 0.01 for the Louvain procedure in order to identify two communities.

### 2 $P_{\mathrm{mc}}$ Proofs and Derivations

#### 2.1 Computation of $P_{\mathrm{mc}}$

To compute the Bayes risk for a general classifier $\delta(\boldsymbol{x})$ under the 0-1 loss, we first evaluate its posterior expected loss $\mathbb{E}_\theta\big[L(\delta(\boldsymbol{x}),\theta) \mid \boldsymbol{x}\big]$ as follows:

$$\begin{aligned}
\mathbb{E}_\theta\big[L(\delta(\boldsymbol{x}),\theta) \mid \boldsymbol{x}\big] &= \Pr(\theta \neq \delta(\boldsymbol{x}) \mid \boldsymbol{x}) \\
&= \sum_{j=1}^{K} \Pr(\delta(\boldsymbol{x}) = j,\, \theta \neq j \mid \boldsymbol{x}) \\
&= \sum_{j=1}^{K} \Pr(\theta \neq j \mid \boldsymbol{x}) \Pr(\delta(\boldsymbol{x}) = j \mid \boldsymbol{x}) \\
&= \sum_{j=1}^{K} \sum_{i \neq j} \Pr(\theta = i \mid \boldsymbol{x}) \Pr(\delta(\boldsymbol{x}) = j \mid \boldsymbol{x}) \\
&= \sum_{j=1}^{k} \sum_{i \neq j} \pi_i(\boldsymbol{x}) \Pr(\delta(\boldsymbol{x}) = j \mid \boldsymbol{x}) \\
&= \sum_{j=1}^{k} (1 - \pi_j(\boldsymbol{x})) \Pr(\delta(\boldsymbol{x}) = j \mid \boldsymbol{x})
\end{aligned}$$

Note that $\Pr(\theta \neq j \mid \boldsymbol{x}, \delta(\boldsymbol{x})) = \Pr(\theta \neq j \mid \boldsymbol{x})$. Subsequently,

$$\begin{aligned}
P_{\mathrm{mc}} &= \mathbb{E}_{\boldsymbol{x}}\Big[\mathbb{E}_\theta\big[L(\delta(\boldsymbol{x}),\theta) \mid \boldsymbol{x}\big]\Big] \\
&= \int \left( \sum_{j=1}^{K} \sum_{i \neq j} \pi_i(\boldsymbol{x}) \Pr(\delta(\boldsymbol{x}) = j \mid \boldsymbol{x}) \right) P(d\boldsymbol{x}) \\
&= \int \left( \sum_{j=1}^{K} (1 - \pi_j(\boldsymbol{x})) \Pr(\delta(\boldsymbol{x}) = j \mid \boldsymbol{x}) \right) P(d\boldsymbol{x})
\end{aligned}$$

#### 2.2 The Cluster Merging Property of $P_{\mathrm{mc}}$

*Proof.* Consider merging two existing clusters $i, j$ to a new combined cluster $k'$. It follows that

$$\alpha_{k'} = \alpha_i + \alpha_j$$

and

$$p(\boldsymbol{x} \mid \theta = k) = \frac{\alpha_i \, p(\boldsymbol{x} \mid \theta = i) + \alpha_j \, p(\boldsymbol{x} \mid \theta = j)}{\alpha_{k'}}$$

Consequently, by applying Bayes rule,

$$\pi_{k'}(\boldsymbol{x}) = \pi_i(\boldsymbol{x}) + \pi_j(\boldsymbol{x}) \tag{1}$$

Let $S$ denote the set of indices of the existing clusters not impacted by the merge, where $|S| = K - 2$. By Eqn (6), $P_{\mathrm{mc}}$ can be written as

$$\begin{aligned} P_{\mathrm{mc}} = 2 &\sum_{m,n \in S,\, m<n} \int \pi_m(\boldsymbol{x}) \pi_n(\boldsymbol{x}) P(d\boldsymbol{x}) \\ &+ 2 \sum_{l \in S} \int \pi_l(\boldsymbol{x}) \pi_i(\boldsymbol{x}) P(d\boldsymbol{x}) + 2 \sum_{l \in S} \int \pi_l(\boldsymbol{x}) \pi_j(\boldsymbol{x}) P(d\boldsymbol{x}) \\ &+ 2 \int \pi_i(\boldsymbol{x}) \pi_j(\boldsymbol{x}) P(d\boldsymbol{x}) \end{aligned}$$

By Eqn (1),

$$2 \sum_{l \in S} \int \pi_l(\boldsymbol{x}) \pi_i(\boldsymbol{x}) P(d\boldsymbol{x}) + 2 \sum_{l \in S} \int \pi_l(\boldsymbol{x}) \pi_j(\boldsymbol{x}) P(d\boldsymbol{x}) = 2 \sum_{l \in S} \int \pi_l(\boldsymbol{x}) \pi_{k'}(\boldsymbol{x}) P(d\boldsymbol{x})$$

and note that,

$$P^{\dagger}_{\mathrm{mc}} = 2 \sum_{m,n \in S,\, m<n} \int \pi_m(\boldsymbol{x}) \pi_n(\boldsymbol{x}) P(d\boldsymbol{x}) + 2 \sum_{l \in S} \int \pi_l(\boldsymbol{x}) \pi_{k'}(\boldsymbol{x}) P(d\boldsymbol{x})$$

It becomes evident that

$$\Delta P^{(i,j)}_{\mathrm{mc}} = P_{\mathrm{mc}} - P^{\dagger}_{\mathrm{mc}} = 2 \int \pi_i(\boldsymbol{x}) \pi_j(\boldsymbol{x}) P(d\boldsymbol{x}) \geq 0 \tag{2}$$

Plugging in the expression of $\Delta P^{(i,j)}_{\mathrm{mc}}$ into Eqn (6) yields

$$P_{\mathrm{mc}} = \sum_{i<j} \Delta P^{(i,j)}_{\mathrm{mc}} \tag{3}$$

□

**Remark** The merging property is specific to the randomized decision rule $\delta_r$ under the 0-1 loss. For the optimal decision rule, $\delta_o$, it can be shown that $P^{\dagger}_{\mathrm{mc}} \leq P_{\mathrm{mc}}$ after merging a pair of existing clusters. However, the quantitive expression for $\Delta P^{(i,j)}_{\mathrm{mc}}$ is analytically intractable.

### 3 Monte Carlo $P_{\mathrm{mc}}$ Comparison

Here we compare $P_{\mathrm{mc}}$ computed using standard cubature methods implemented in the `cubature` R package [2] to $\hat{P_{\mathrm{mc}}}$ estimated using a Monte Carlo (MC) integral (as described in Section 3.1.6). We use 50 replicates to obtain the timing measurements and quantify the uncertainty in the MC integral. We compare the methods in terms of their average time to evaluate $P_{\mathrm{mc}}$ for a given cluster density configuration. The configuration in question consists of three Gaussian clusters with $\pi_k = 1/3$ in $\mathbb{R}^p$ with $I_p$ variance. The cluster are centered at $\mathbf{0}^p$ and $\pm\mathbf{d}^p$, where $\mathbf{d}^p$ is the p-dimensional vector whose elements are all $d$. $d$ is set so that the Euclidean distance between $\mathbf{d}^p$ and the origin is fixed to be 3; i.e. $d = \sqrt{3^2/p}$. This is so that the value of $P_{\mathrm{mc}}$ remains fixed across dimensions and we do not need to worry about the dimensionality affecting the true value of $P_{\mathrm{mc}}$. For the MC integration procedure we use $M = 10^5$ sample points. The results for dimension $p = 1, \ldots, 5$ are presented in Supplementary Table 2 below.

Both approaches produce highly similar values of $P_{\mathrm{mc}}$ across dimensions. The standard deviation of the MC estimates is quite low, indicating stability in the estimation procedure. Additionally, while the cubature method evaluation time rapidly increases in $p$, the time for the MC procedure is roughly constant. These together highlight the MC estimation procedure as a viable and accurate approach to estimate $P_{\mathrm{mc}}$, especially in moderate dimensional data where cubature methods may struggle to reach a solution in a reasonable time.

| D | $P_{\mathrm{mc}}$ | Elapsed (s) | $\hat{P_{\mathrm{mc}}}$ | $\sigma(\hat{P_{\mathrm{mc}}})$ | Elapsed (s) |
|---|---|---|---|---|---|
| 1 | 0.13144 | 0.00302 | 0.13143 | 0.00047 | 0.76102 |
| 2 | 0.13144 | 0.04638 | 0.13139 | 0.00050 | 0.69586 |
| 3 | 0.13144 | 1.16948 | 0.13136 | 0.00048 | 0.68714 |
| 4 | 0.13144 | 7.58432 | 0.13136 | 0.00046 | 0.73066 |
| 5 | 0.13145 | 8.61848 | 0.13153 | 0.00040 | 0.72620 |

Supplementary Table 2: $P_{\mathrm{mc}}$ values computed using cubature methods and Monte Carlo integration based on 50 replicates. The $P_{\mathrm{mc}}$ and $\hat{P_{\mathrm{mc}}}$ values are averages across all replicates. $\sigma(\hat{P_{\mathrm{mc}}})$ is the standard deviation of the MC $P_{\mathrm{mc}}$ estimate across the 50 replicates. Elapsed time (in seconds) is the average time for a single replicate.

### 4 PHM Heatmap Spline-based Scaling

In order to produce a $P_{\mathrm{mc}}$ scale more indicative of the distance between two clusters, we consider two univariate Gaussian clusters $N(-\mu/2, 1)$ and $N(\mu/2, 1)$ with equal mixing proportions such that the distance between the cluster centers is $\mu$. We first compute $P_{\mathrm{mc}}$ for values of $\mu \in \{0.1, 0.2, \ldots, 50\}$ using standard quadrature methods. We then fit a cubic spline using the $\log_{10} P_{\mathrm{mc}}$ values to predict $\mu$ as a way to transform from the $P_{\mathrm{mc}}$ scale to a linear distance scale ($s(x)$).

Figure 21 shows a comparison of different scaling approaches: fitting a spline directly to the $P_{\mathrm{mc}}$ values ($s(P_{\mathrm{mc}})$), using the $-\log_{P_{\mathrm{mc}}}$ values directly, and fitting a spline to the $\log_{10} P_{\mathrm{mc}}$ values ($s(\log_{10} P_{\mathrm{mc}})$). In this illustration, we use $\mu \leq 30$ to fit the splines and try to predict the distances for $\mu > 30$. Directly using the log scaling appears to distort the distance measure significantly, as smaller distances are under-estimated whereas large distances are inflated. Using the spline to estimate the distance directly from $P_{\mathrm{mc}}$ appears to be predicting roughly constant values outside of the training data range. The spline fit to the $\log_{10} P_{\mathrm{mc}}$ values appears to fit the data properly, preserving the linear relationship with distance in the testing data.

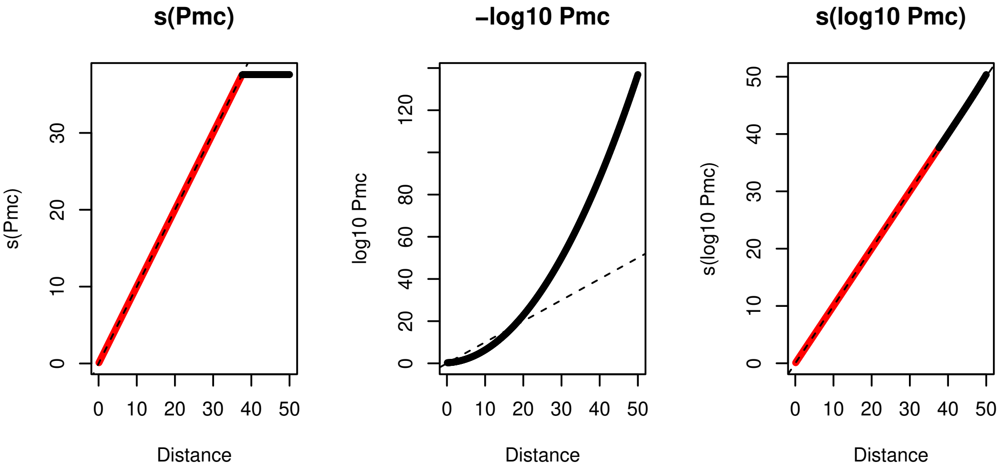


Supplementary Figure 21: **Comparison of $P_{\mathrm{mc}}$ distance scaling options** Comparison of predicted "distance" measures from *(Left)* a spline fit directly to $P_{\mathrm{mc}}$ values, *(Center)* the $-\log_{10} P_{\mathrm{mc}}$ values, and *(Right)* a spline fit to the $\log_{10} P_{\mathrm{mc}}$ values. For the spline-based scalings, red points indicate those used to fit the spline, black points are predicted from the corresponding $P_{\mathrm{mc}}$ values. Dashed line indicates $y = x$.

[2] Balasubramanian Narasimhan, Steven G. Johnson, Thomas Hahn, Annie Bouvier, and Kiên Kiêu. *cubature: Adaptive Multivariate Integration over Hypercubes*, 2024. R package version 2.1.1.
